# Engineering *Agrobacterium tumefaciens* adhesion to target cells

**DOI:** 10.1101/2022.02.15.480547

**Authors:** Xavier Pierrat, Alix Pham, Jeremy P. H. Wong, Zainebe Al-Mayyah, Alexandre Persat

## Abstract

*Agrobacterium tumefaciens* is a plant pathogen commonly repurposed for genetic modification of crops. Despite its versatility, it remains inefficient at transferring DNA to many hosts, including to animal cells. Like many pathogens, physical contact between *A. tumefaciens* and host cells promotes infection efficacy. Thus, improving the strength and specificity of *A. tumefaciens* to target cells has the potential for enhancing DNA transfer for biotechnological and therapeutic purposes. Here we demonstrate a methodology for engineering genetically-encoded exogeneous adhesins at the surface of *A. tumefaciens*. We identified an autotransporter gene we named Aat, that is predicted to show canonical β-barrel and passenger domains. We engineered the β-barrel scaffold and linker (Aat_β_) to display synthetic adhesins susceptible to rewire *A. tumefaciens* to alternative host targets. As a proof of concept, we leveraged the versatility of a VHH domain to rewire *A. tumefaciens* adhesion to yeast and mammalian hosts displaying a GFP target receptor. Finally, to demonstrate how synthetic *A. tumefaciens* adhesion can improve transfer to host cells, we showed improved protein translocation into HeLa cells using a sensitive split luciferase reporter system. Engineering *A. tumefaciens* adhesion has therefore a strong potential in generating complex heterogeneous cellular assemblies and in improving DNA transfer efficiency against non-natural hosts.

## Introduction

*A. tumefaciens* is a soil bacterium able to transfer DNA fragments tens of kilobases-long to plant host cells. In its natural environment, *A. tumefaciens* transfers a pathogenic DNA fragment called transferred-DNA (T-DNA). *A. tumefaciens* has been repurposed into a potent gene delivery tool for a broad range of plants, yeasts and fungi^1–3^. Using disarmed strains lacking pathogenic T-DNA, plant engineers introduce new genes in target organisms such as maize, rice and wheat^4–6^. Genetic modifications include random integration or targeted editing using zinc-finger nucleases or CRISPR/Cas9 ^4^.

*A. tumefaciens* uses the type IV secretion (T4SS) system to inject DNA into host cells. The T4SS functions upon host surface contact, so that attachment to target cells promotes T-DNA transfer^7–9^. For synthetic application, forcing cell-cell interaction, for instance by wounding the plant or by using syringe-or vacuum-driven agroinfiltration, is usually recommended^10^. The addition of extracellular cellulose increases bacterial adhesion to improve T-DNA transfer efficiency to recalcitrant plant cells^11^. Altogether, these studies suggest that increased bacterial adhesion favors T-DNA transfer. However, our understanding of how *A. tumefaciens* adheres to host plant surfaces remains incomplete^8,9^. Despite this knowledge gap, engineering adhesion has the potential to improve gene delivery to plants or alternative targets, for example by broadening the host range of *A. tumefaciens*.

In Gram-negative bacteria, outer-membrane proteins called autotransporters help display functional proteins moieties at the cell surface, including adhesins^12^. Autotransporters are promising candidate scaffolds for surface display of exogenous proteins^13^. They belong to the type V secretion system (T5SS) family, and are composed of a β-barrel domain fused to a passenger domain with an α-helical linker^14^. The β-barrel anchors the autotransporter to the outer membrane. The passenger domain translocates by traversing the β-barrel, thereby exposing itself at the cell surface and conferring function. In contrast to other secretion systems, autotransporters are expressed as a single protein. This allows for substitution of the passenger domain to other peptides as a way to switch autotransporter function. For example, the *Pseudomonas aeruginosa* EstA autotransporter has been used to display a variety of lipases, with applications in whole-cell biocatalysis and screening of enzyme libraries^14^.

Adhesion plays a crucial function in host microbe interactions, which can be repurposed for synthetic applications. Fusing an alternative receptor to an autotransporter is a convenient way to engineer bacterial adhesion to a specific host target. For example, fusions of non-natural receptors to the *E. coli* intimin autotransporter efficiently rewires adhesion to a wide range of host cells. Specifically, antigen-binding domain from camelid heavy-chain antibodies (<u>v</u>ariable <u>h</u>eavy chain of <u>h</u>eavy-chain antibodies, VHH) fused to intimin provide adhesion properties to target ligands^15^. Utilizing VHH has many advantages as they are single-chained, short (less than 130 amino acids-long) and have a robust structure^16^. Moreover, they are more efficiently displayed at the surface of bacteria than human single-chain fragment variables (scFv)^17^. Finally, screening for VHH forms with affinity to new targets is streamlined, for example via phage display starting from naïve DNA libraries^18^. Using the intimin-VHH display system, Fraile *et al*. targeted *P. putida* to abiotic surface coated with the target antigen^19^. Salema *et al*. screened libraries against cancer biomarkers by selecting bacteria binding to live mammalian cells^20^. Glass *et al*. created an adhesin toolbox that enables the assembly of complex *E. coli* communities^21^. Finally, Ting *et al*. performed targeted bacterial killing using programmed adhesion that directs toxin injection^22^. Thus, synthetic adhesion can confer modularity and specificity in assembling bacterial consortia or attaching to alternative hosts^23,24^.

Here, we aimed at engineering *A. tumefaciens* adhesion to improve protein delivery to alternative targets, with the ultimate goal to program DNA transfer into alternative hosts. While using an intimin scaffold is intuitive, it is not a practical solution. Intimin has a topology belonging to the reverse autotransporter family, also called type Ve secretion system^25–27^. *A. tumefaciens* C58 does not possess any annotated reverse autotransporter, and consequently likely does not express the right variants of the associated chaperones that enable proper insertion in the outer membrane and/or translocation of the passenger domain^25^. Therefore, we explored alternative autotransporters in order to engineer synthetic adhesins at the surface of *A. tumefaciens*.

Here, we engineered a modular synthetic adhesin display system for *A. tumefaciens*. To achieve this, we investigated the putative autotransporter Atu5364 which we renamed Aat (as *<u>A</u>grobacterium* <u>a</u>utotransporter) as candidate scaffold for adhesin display. We successfully repurposed Aat to display multiple functional receptors targeting a variety of ligands, including VHH, lectins and arginine-glycine-aspartic acid (RGD) peptide. We then demonstrated that displaying a nanobody-based VHH anti-GFP by Aat fusion strongly improved *A. tumefaciens* attachment to alternative hosts including yeast and mammalian cells displaying GFP. Finally, and as a proof of concept, we provide preliminary evidence that synthetic display of VHH improves transfer of type IV secretion system proteins to mammalian host cells.

## Results

### Identification of an A. tumefaciens autotransporter scaffold

The genome of *A. tumefaciens* is annotated with two uncharacterized autotransporters of the type Va secretion system (T5aSS), namely *atu5354* and *atu5364*, both located on the cryptic megaplasmid pAtC58. We only found a signal peptide encoded within the sequence of *atu5364*, but not in *atu5354*, which we therefore disregarded as a potential candidate scaffold (Figure S1). We then modeled *<u>A</u>grobacterium* <u>a</u>utotransporter Atu5364 (Aat) using RoseTTaFold and obtained an estimate of its three dimensional conformation (Figure 1A)^28,29^. The predicted structure shows a C-terminal scaffold (β-barrel and α-helix) holding a passenger formed of repeated parallel β-strand repeats organized in a β-solenoid, a prevalent structure in T5aSS autotransporters^30^. Aat harbors a proline-rich linker between the scaffold and the folded region of the passenger domain. A potential function for this linker is to remain unfolded and extend to help the passenger domain reach distant targets, for example in attachment^31^. We did not identify a clear homolog for the passenger domain, but our analysis points toward a function in adhesion (Table S1)^32–37^. Altogether, the predicted scaffold, signal peptide and long linker make Aat a promising candidate for the autotransporter-based display of non-natural passenger domains.

**Figure 1:**
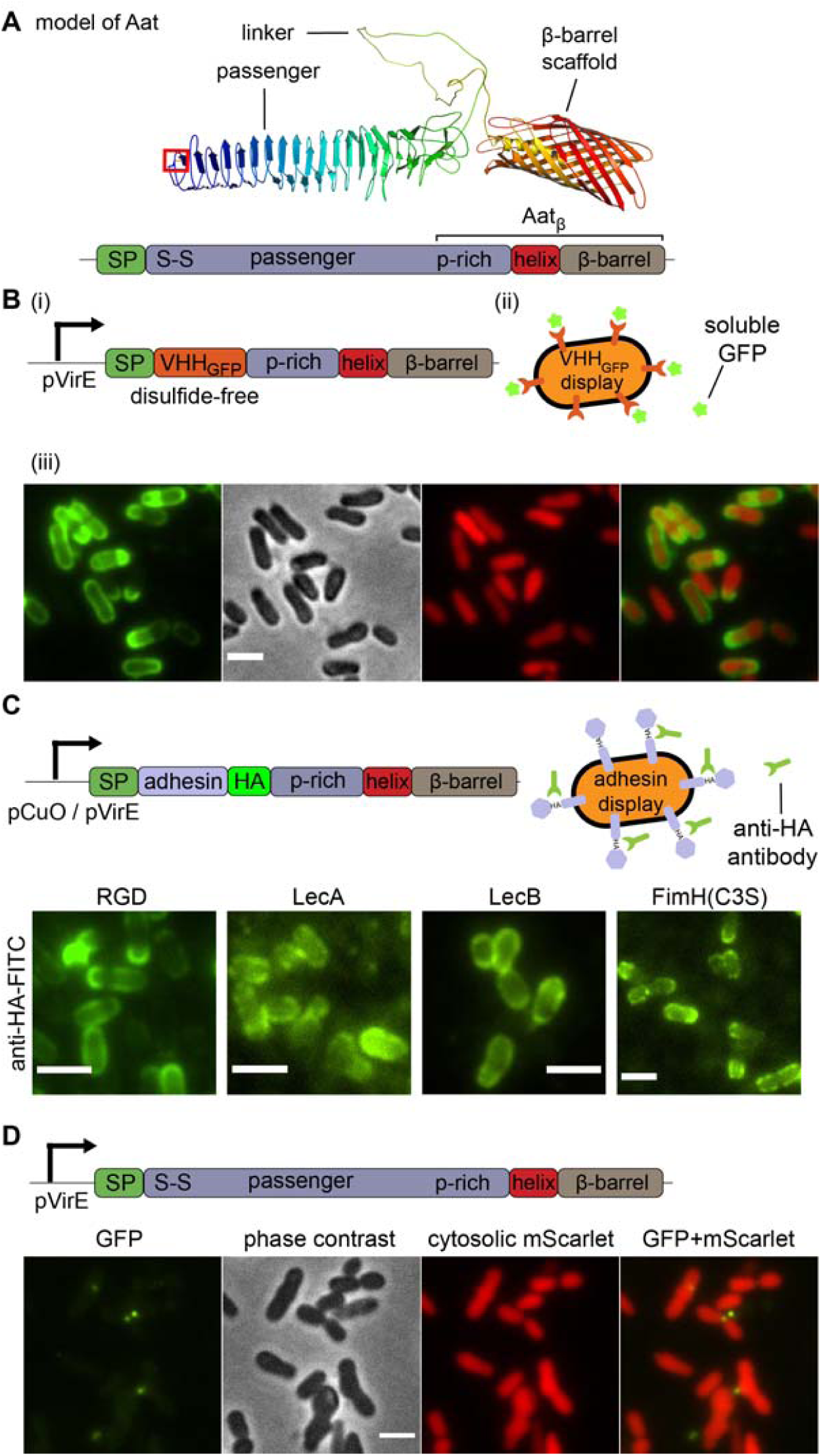
Protein display in *A. tumefaciens* by fusion to Aat_β_. (A) Structure of Aat modeled by RoseTTaFold. Aat is a T5aSS autotransporter with a C-terminal β-barrel inserted at the outer membrane. A passenger domain consists of β-strands capped by a disulfide bond (C35-C48, S-S, framed in red). Aat_β_ is composed of the β-barrel, α-helix and proline-rich linker (p-rich). SP = signal peptide. (B) Fusion of cysteine-free VHH anti-GFP (VHH_GFP_) to Aat_β_ is displayed at the surface of *A. tumefaciens*. (i) Diagram of the construct placed under a VirE promoter (pVirE). (ii) Illustration of *A. tumefaciens* expressing VHH_GFP__Aat_β_ (red) binding to GFP (green stars). (iii) *A. tumefaciens* expressing VHH_GFP__Aat_β_ binds diffusible GFP at its surface. Cytosolic mScarlet highlights the surface GFP signal. (C) Display of arginine-glycine-aspartate (RGD), *P. aeruginosa* LecA, LecB and disulfide-free *E. coli* FimH at the surface of *A. tumefaciens*. The constructs are placed under a pVirE or a synthetic cumic acid-inducible promoter (pCuO) with an HA-tag allowing for visualization. Bacteria were stained with anti-HA antibody conjugated with fluorescein (FITC). (D) Negative control: overexpression of Aat wild-type under a VirE promoter, followed by staining with GFP. Bacteria have aberrant shapes and are not stained with GFP. Bars, 2 µm.

### Protein display in A. tumefaciens

We then moved on to use Aat as a scaffold for protein display in *A. tumefaciens*. We first fused an HA-tag between the endogenous passenger domain and the linker domain. We expressed the fusion under a cumic acid-inducible promoter to decouple expression from virulence^38^. After staining with FITC-conjugated anti-HA antibodies, these cells showed low expression with patchy localization at the cell surface (Figure S2B). We noticed that the native passenger of Aat contains a disulfide bond located at the N-terminal end of the native passenger domain, which may prevent translocation (Figure 1A)^39^. We generated a cysteine-free mutant version of the passenger domain with the HA-tag fusion. Staining and microscopy showed that expression levels of the cysteine-free mutant were much stronger than wild-type (WT) and that the localization was uniform along the bacterium surface (Figure S2C).

To demonstrate the capacity of Aat in displaying alternative functional passenger domains, we fused a VHH domain targeting GFP to the Aat β-barrel scaffold with its proline-rich linker (which we call Aat_β_). We initially screened a variety of fusion strategies, which consisted in either replacing parts of the passenger domain (amino acids 35-160, 35-512 or 161-512) or in introducing VHH in front, within or after the folded domain of the passenger (at C35, D161 or A513) (Supplementary figure S3A). The constructs were expressed from binary vectors under the control of the pVirE promoter (Supplementary table 2)^40^. Incubating bacteria expressing these constructs with GFP shows that the scaffold is properly folded and inserted in the outer membrane. However, these constructs caused frequent cell death (Figure S3B, C). We hypothesize that VHH is folded but accumulates in the periplasm, causing toxicity.

We noticed that VHH contains a disulfide bond that again could prevent passenger domain translocation^39^. Mutating cysteines from VHH anti-GFP has little effect on affinity^41^. We therefore cloned a disulfide-free VHH by site-directed mutagenesis of C24A and C98V. We then displayed the disulfide-free VHH anti-GFP (VHH_GFP_) by fusion to Aat_β_ (VHH_GFP__Aat_β_) in *A. tumefaciens* and compared it to the native VHH. We expressed this construct in a strain constitutively expressing cytoplasmic mScarlet as viability reporter. After expressing and staining VHH_GFP__Aat_β_ with GFP, we observed enhanced fluorescent signal at the bacterium surface compared to the native, disulfide-containing version, while not affecting bacterial viability (Figure 1B, Figure S3C, D). Of note, the distribution of the GFP signal seemed more concentrated at the narrower side of the bacteria, which might be due to intrinsic, yet unknown, properties of the Aat_β_ scaffold. In summary, Aat_β_ efficiently displays functional VHH_GFP_ at the surface of live bacteria.

To illustrate the modularity of the display scaffold, we sought to display alternative receptors. We turned to the arginine-glycine-aspartate (RGD) motif which targets integrin ligands, and two disulfide-free lectins from *Pseudomonas aeruginosa*, namely LecA and LecB which respectively bind to galactose and fucose^42–45^. We fused each of the receptors to Aat_β_. In each, we also inserted an HA-tag between the adhesin and Aat_β_ for characterization of display. Upon induction and staining with a green fluorescent anti-HA antibody, all constructs showed fluorescence at their periphery. All maintained WT levels of a constitutively-expressed cytosolic mScarlet demonstrating cell viability (Figures 1C, S4A-E). To further demonstrate the process of engineering adhesins for Aat_β_-based display, we displayed the FimH fimbrial tip of uropathogenic *E. coli*. FimH harbors a disulfide bond between C3 and C44. Mutations of one of FimH cysteines to a serine does not affect affinity to mannose at no-and low shear rates^46^. Hence, we generated a FimH(C3S) mutant that is deficient in forming disulfide bonds and compared the display efficiency with the WT. Like for VHH display, the removal of the disulfide bond greatly improved display (Figures 1C, right panel and S4F and G). Altogether, this shows that Aat_β_ is highly modular for passenger display and likely can display a wide variety of short peptides, tags, and other disulfide-free β-stranded proteins. For clarity and demonstration purposes, we subsequently focused on the display of VHH_GFP_ adhesin as it enables quantitative control at the adhesin level, but also at the host ligand level.

To improve the genetic stability of the constructs, we generated markerless genomic insertion of VHH_GFP__Aat_β_ and compared it to plasmid-based expression. More specifically, we integrated VHH_GFP__Aat_β_ at the *virE2* locus on the Tumor-inducing (Ti) megaplasmid (Supplementary figure S5). We next induced bacteria with increasing concentrations of inducer (acetosyringone titration) in both of the plasmid-borne and chromosomally-integrated version. We stained the bacteria with recombinant GFP to quantify the display efficiency of VHH_GFP__Aat_β_ (Figure 2). Both Ti plasmid-integrated and binary vector-based displayed VHH_GFP_ at high and very high levels, a small difference probably likely due to plasmid copy number. In conclusion, both binary vector-based and Ti plasmid-integrated versions are compatible with VHH_GFP_ display.

**Figure 2:**
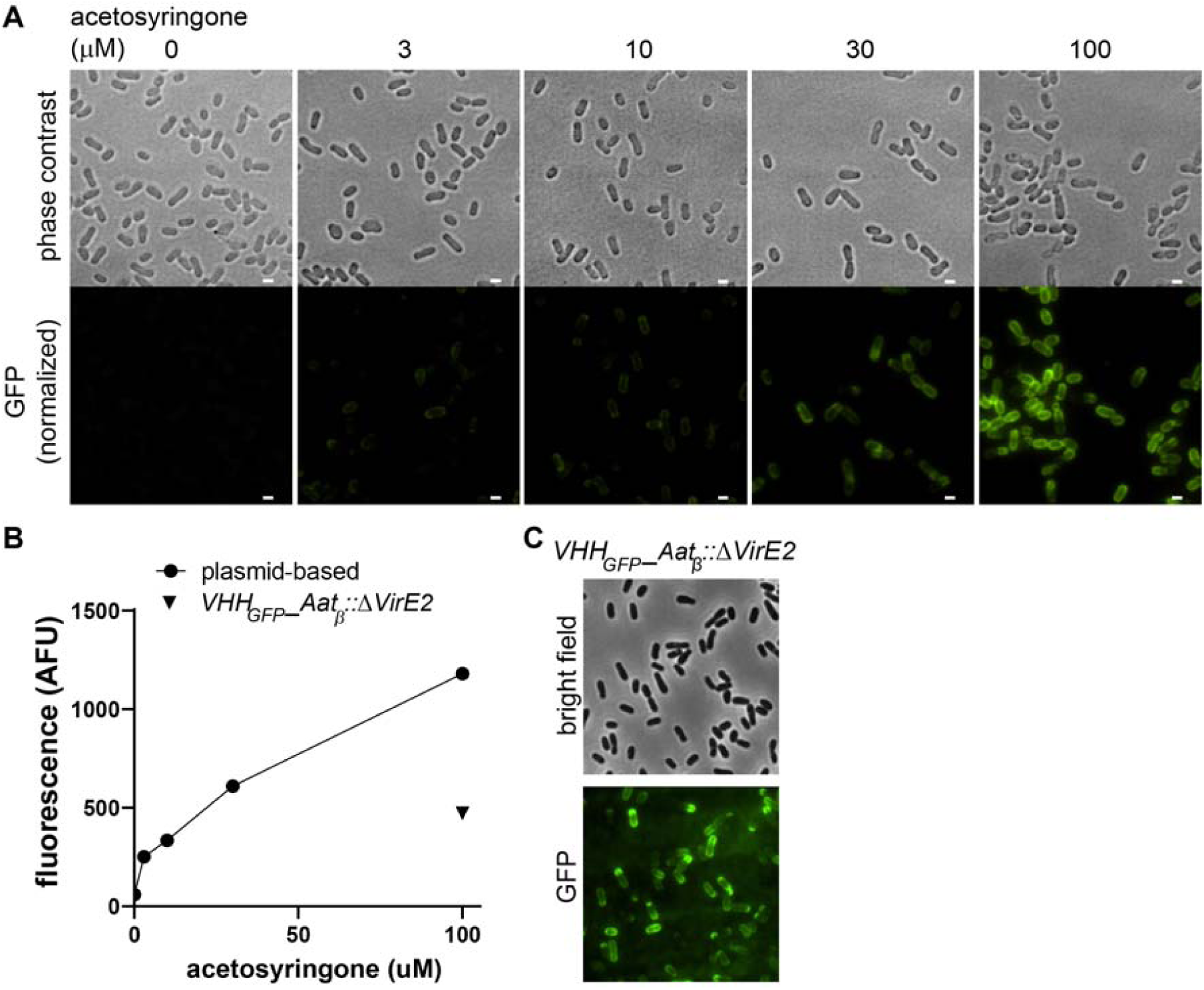
pVirE is a potent inducible promoter for VHH_GFP__Aat_β_ on binary vector and Tumor-inducing megaplasmid. (A) Acetosyringone induction of VHH_GFP__Aat_β_ from a VirE promoter on a binary vector. *A. tumefaciens* transformed with pVirE – VHH_GFP__Aat_β_ (pXP213, table S2) was induced with increasing concentrations of acetosyringone and stained with recombinant eGFP. GFP fluorescence intensity scale is identical between images. (B) Mean bacterial GFP intensity as a function of inducer concentration. (C) VHH_GFP_*_*Aat_β_ can also be expressed from a stable insertion in the disarmed tumor-inducing plasmid (*A. tumefaciens VHH*_*GFP*_*_Aat*_β_::Δ*VirE2*, see table S3). Bars, 1 µm.

### Attachment of engineered A. tumefaciens to yeast

Synthetic adhesin display opens the possibility of engineering bacteria with emerging functions, such as the assembly of complex multicellular structures, the destruction of cancer cells, or can be applied in investigations of adhesion to host cells^21,47^. With these applications in mind, we explored the potential of *A. tumefaciens* with synthetic Aat_β_ display in providing attachment to non-plant hosts. Forcing bacteria-host contact improves Agrobacterium-mediated transformation of yeast cells^1,48^. We therefore first focused on testing whether the synthetic VHH_GFP__Aat_β_ construct promotes attachment to yeast cells. By fusing GFP to cell-wall-anchoring proteins, we constructed a GFP-displaying *S. cerevisiae* in the strain eby100, commonly used in yeast display libraries (Figure 3A)^49^. Then, we separately induced GFP-display in yeast and expression of VHH_GFP__Aat_β_ in *A. tumefaciens*. We mixed the two strains and imaged the consortium by confocal microscopy. Confocal sections showed that bacteria displaying VHH_GFP_ strongly bound to the yeast cell wall, to the extent of imprinting their shape into the cell walls (Figure 3B). We then compared the binding to GFP-displaying yeast in the absence of VHH_GFP__Aat_β_ and observed a decrease in the number of bacteria bound per yeast cell (Figure 3C). In order to unambiguously attribute binding to the specificity of VHH_GFP_ to GFP, we compared this to GFP-negative yeast and *A. tumefaciens* displaying a cysteine-free VHH anti-mCherry (VHH_mCherry_). The number of bacteria attached per yeast cell only increased when the VHH_GFP__Aat_β_ construct was expressed and when yeast displayed GFP (Figure 3D). This shows that our scaffold can rewire *A. tumefaciens* to the yeast cells, opening the possibility of generating complex interkingdom assemblies. The low binding of *A. tumefaciens* displaying VHH_mCherry_ additionally demonstrates the specificity of synthetic adhesion.

**Figure 3:**
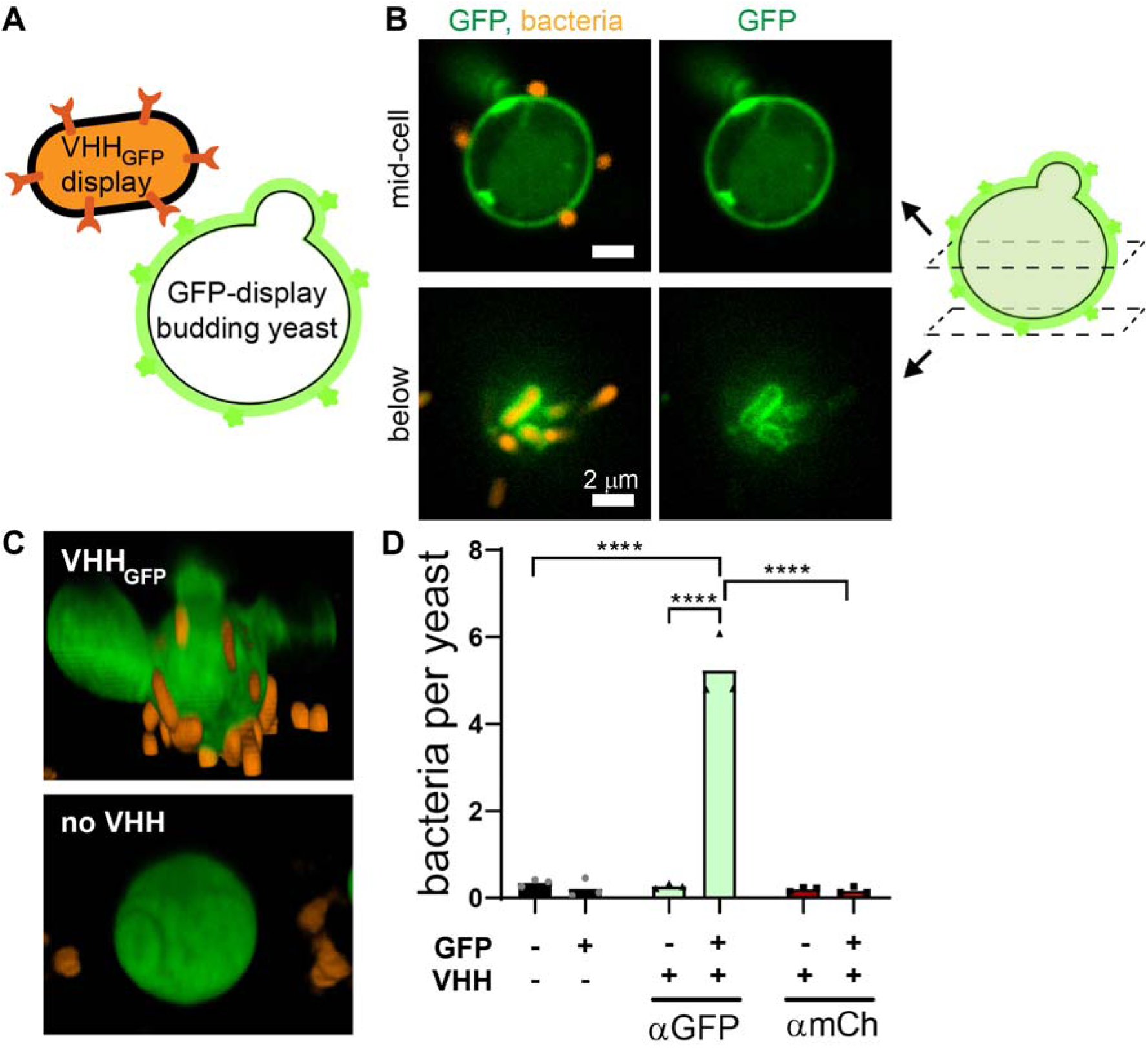
Synthetic adhesion of *A. tumefaciens* VHH_GFP__Aat_β_ to GFP-displaying yeast cells. *A. tumefaciens mScarlet* VHH_GFP__Aat_β_ binds to GFP displayed at the surface of *S. cerevisiae*. Representative confocal images of *A. tumefaciens mScarlet* VHH_GFP__Aat_β_ attaching to GFP-displaying yeast cell. In the lower panels, bacterial shape imprints are visible. (C) 3D visualization of *A. tumefaciens mScarlet* (orange) binding to *S. cerevisiae* displaying GFP (green) in the absence or presence of VHH_GFP_ display. (D) Number of *A. tumefaciens mScarlet* cells attached per yeast cell [no VHH: empty vector pFGL815, αGFP: pVirE - VHH_GFP__Aat_β_, αmCh: pVirE - cysteine-free VHH anti-mCherry (VHH_mCherry__Aat_β_), see table S2). Bars represent the mean of biological triplicates. Statistical tests: two-way ANOVA and Sidak *post-hoc* test (****, P<10^−4^).

### Attachment of engineered A. tumefaciens to mammalian cells

To demonstrate the capability of the synthetic Aat_β_-based display for therapeutic purposes, we tested how engineered *A. tumefaciens* could specifically bind human cells. To achieve this, we employed HeLa cells displaying GFP at their surface by fusion to a CD80 transmembrane domain^47^. We incubated HeLa GFP-display with bare or displaying VHH_GFP_ *A. tumefaciens*. After washing of unbound bacteria, we acquired confocal images to quantify the average number of bacteria per HeLa cell. We observed an increase in the number of bound bacteria when HeLa displayed GFP and *A. tumefaciens* displayed VHH_GFP_ simultaneously. However, there was no increase in binding when only the receptor or its ligand were expressed (Figure 4Ai-iii and B). By preincubating *A. tumefaciens* with GFP, we were able to prevent binding to host cells, demonstrating specificity and the potential for external inhibition of adhesion (Figure 4iv and B). In addition, *A. tumefaciens* displaying VHH_mCherry_ could not attach to HeLa displaying GFP, further demonstrating specificity (Figure 4Av and B). These results held across cells lines, as *A. tumefaciens* attached in a similar manner to GFP-displaying HEK293T cells (Figure S6).

**Figure 4:**
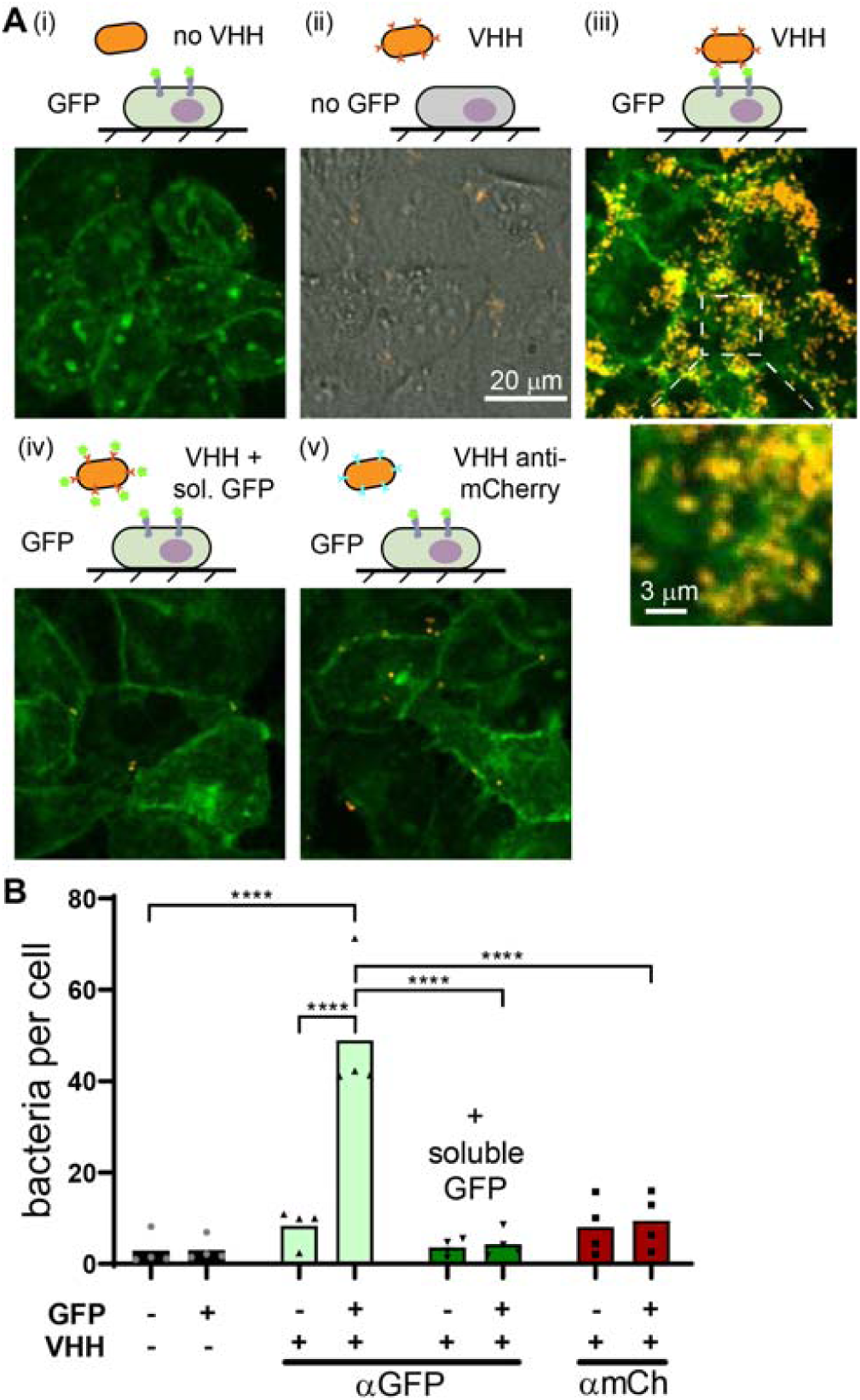
Synthetic adhesins promote specific *A. tumefaciens* binding to HeLa cells. (A) Maximum intensity projections of *A. tumefaciens mScarlet* binding to HeLa cells. (i) no VHH_GFP__Aat_β_ and GFP-display, (ii) VHH_GFP__Aat_β_ and no GFP-display, (iii) VHH_GFP__Aat_β_ and GFP display, (iv) VHH_GFP__Aat_β_ and GFP-display prevented by soluble GFP and (v) VHH_mCherry__Aat_β_ and GFP display. All images are overlays of GFP (green) and bacteria (orange) except (ii) where we use brightfield to visualize non-fluorescent HeLa. (B) Average number of bacteria per HeLa cells. HeLa were uninduced (GFP -) or induced for GFP display (GFP +), *A. tumefaciens mScarlet* was transformed with an empty vector (VHH -), pVirE – VHH_GFP__Aat_β_ (αGFP) or pVirE – VHH_mCherry__Aat_β_ (αmCh). Dark green set of bars show conditions where soluble eGFP was added to saturate VHH_GFP_ receptors. Each symbol is a biological replicate, bars represent the means of four biological replicates. Statistical tests: two-way ANOVA and Sidak post hoc test (****, P<10^−4^).

### Monitoring VirE2 transfer using split NanoLuc

We next aimed at targeted delivery into mammalian cells using synthetic adhesion. Helper proteins VirE2 and VirD2 increase T-DNA transfer efficiency in experiments involving artificial T-DNA transfection^50^. Following binding to the target cell and T4SS assembly, VirD4 actively recruits helpers proteins such as VirD2 and VirE2 to the T4SS machinery for injection into the target cell’s cytosol. Consequently, the cytosol of the target cell is the first compartment through which the T-DNA and helper proteins navigate.

To sensitively monitor protein transfer, we employed a split luciferase (NanoLuc) approach where the translocated protein is fused to one fragment (HiBit) and the host cell expresses a complementary fragment (LgBit). Successful injection into the host leads to NanoLuc complementation and hence to an increase in luminescence signal from the host^51–53^. VirE2 is the most abundant Vir protein and is often fused to the short fragment of split fluorescent proteins by insertion into an internal permissive site^54,55^. We therefore generated an *A. tumefaciens* strain expressing a HiBit::VirE2 internal fusion under the VirE promoter (Figure 5A). As reporter cell line, we further engineered the GFP-displaying HeLa cell line to constitutively express the LgBit fragment. We controlled LgBit complementation by transiently transfecting cells with a plasmid driving HiBit::VirE2 expression. We measured a strong increase in luminescence signal, suggesting that HiBit internally fused to VirE2 efficiently complements LgBit (Figure S7).

**Figure 5:**
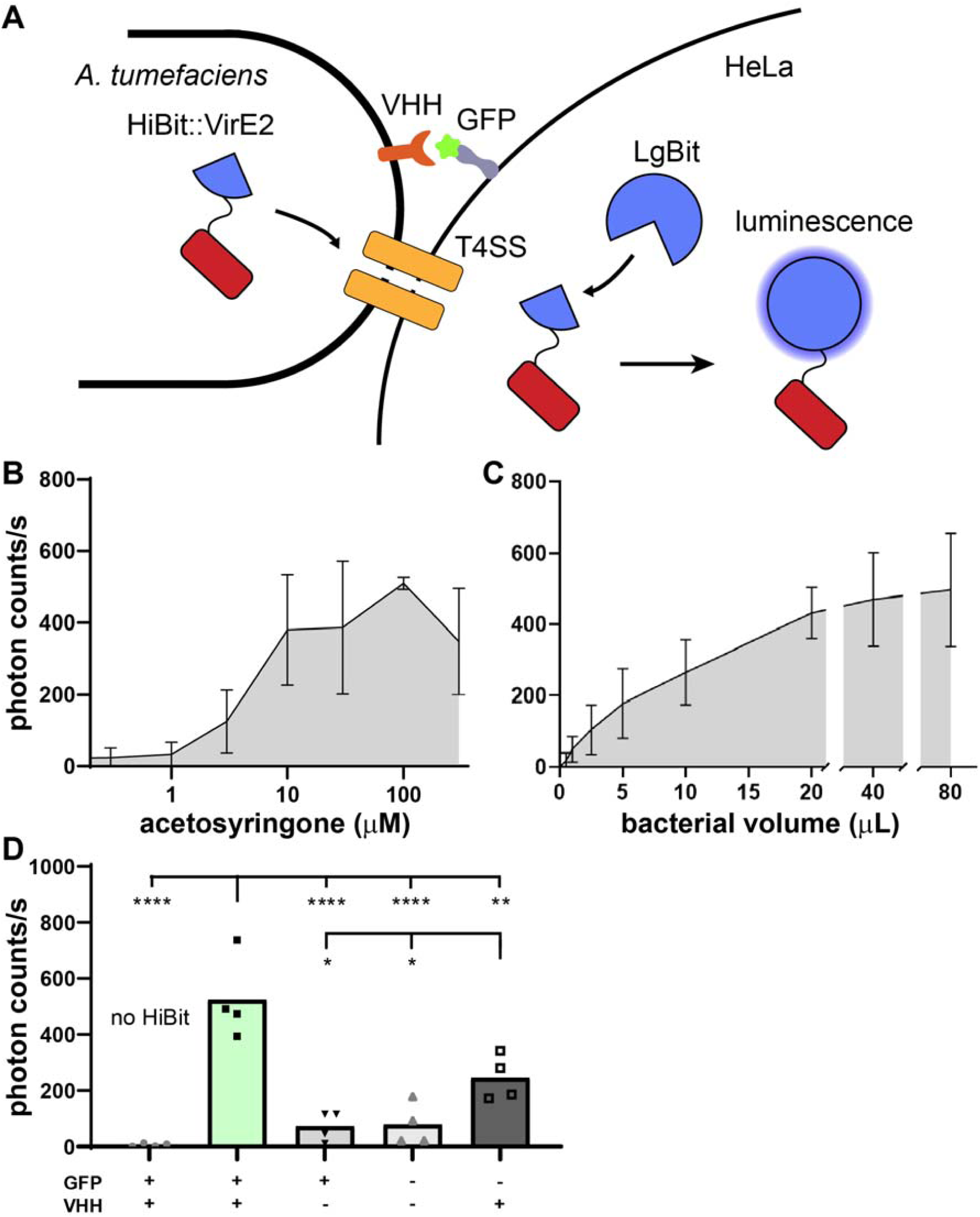
VHH_GFP__Aat_β_ improves VirE2 transfer to HeLa cells. (A) Overview of the split luciferase (NanoLuc) system: NanoLuc is split in two fragments, LgBit and HiBit. *A. tumefaciens VHH*_*GFP*_*_Aat*_β_::Δ*VirE2* expresses an internal fusion of HiBit into VirE2. The target host cell expresses LgBit. Upon successful VirE2 transfer, NanoLuc activity should be restored. (B) HiBit::VirE2 expressing bacteria were induced with increasing concentrations of acetosyringone and added to GFP-displaying HeLa cells expressing LgBit. (C) VirE2 transfer as a function of bacterial load, controlled by volume of culture added to GFP-displaying HeLa cells expressing LgBit. 100 µM inducer concentration. (B and C) Error bars represent the standard deviation of biological triplicates. (D) VHH_GFP__Aat_β_ promotes VirE2 transfer in a GFP-display dependent manner. Bacteria were induced with 100 µM acetosyringone and added to LgBit-expressing reporter HeLa cells with or without GFP ligand at their surface. Each bar represents the means of biological triplicates. Statistical tests: one-way ANOVA followed by Dunnett’s post hoc test (**** P<10^−4^, ** P<0.01, * P<0.05).

We next challenged HeLa GFP display expressing LgBit with *A. tumefaciens* expressing VHH_GFP__Aat_β_ and HiBit::VirE2. We first modulated either inducer concentration or multiplicity of infection. Luciferase signal intensity increased with induction levels at constant multiplicity of infection (Figure 5B), and with multiplicity of infection at constant induction (Figure 5C). This implies that increased in bacterial virulence or increased in number of bacteria both increase VirE2 transfer to HeLa cells.

We then investigated the specificity of VirE2 transfer to the synthetic binding. Based on the titrations (Figure 5B, C), we infected HeLa cells with 22 µL of bacterial culture induced with 100 µM acetosyringone. We first verified that the signal was specific to HiBit::VirE2 translocation by measuring luminescence of a native VirE2 in the same system. We measured only background luminescence levels, order of magnitude smaller than for the HiBit::VirE2 fusion (Figure 5D). In addition, infecting HeLa with or without GFP display with an *A. tumefaciens* strain lacking VHH adhesin display but expressing HiBit::VirE2 abolished luminescence. These experiments demonstrate that VHH display is required for the increase in VirE2 transfer. We however found one discrepancy: luminescence was still high—while statistically significantly lower than when both adhesins are present—when VirE2 is transferred to HeLa without expression of GFP receptors (Figure 5D). To explain this discrepancy, we checked for the possibility of leaky GFP expression. We inhibited binding by pre-incubating *A. tumefaciens* VHH with recombinant GFP or VHH to block adhesion from bacterial or host side respectively. We included a control by adding recombinant mCherry, which should not interfere with VHH in binding GFP on host cells. Luminescence decreased about 2-fold compared to non-induced GFP-display, still partly explaining the GFP-unspecific signal increase (Figure S8A).

We attribute the remaining signal to VirE2 transfer independent of adhesion through an unknown mechanism. We therefore investigated the impact of the Aat_β_ scaffold on transfer. We displayed either a disulfide-free VHH anti-mCherry, or an empty passenger domain, and repeated the experiment. In the absence of specific adhesion to GFP, we observed a similar VirE2 transfer as in the absence of GFP-display (Figure S8B, second to last columns). This suggests that Aat_β_ has an intrinsic contribution to VirE2 transfer. We next verified if this intrinsic contribution of Aat_β_ occurs before contact (e.g. lysis) or upon contact. By comparing the signal of reporter cells onto which we applied either overnight supernatant, resuspended bacteria, we observed that the signal increase was due to VirE2 translocation from bacteria but not VirE2 secreted in the supernatant (Figure S8C). However, lysate of *A. tumefaciens* expressing VHH_GFP__Aat_β_ and HiBit::VirE2 showed a strong increase in luminescence when incubated with HeLa GFP (Figure S8C), which suggests that mammalian cells actively uptake VirE2, by a mechanism that remains to be elucidated. One unifying hypothesis about the non-specific fraction of VirE2 transfer would be that overexpression of Aat_β_ slightly increases unspecific binding to target cells (Figure 4B), reaching a sufficient threshold for VirE2 transfer, disregard of the specificity of the VHH (Figures 5D and S8B). In conclusion, we demonstrated that synthetic adhesion promotes VirE2 transfer to host cells, a process which is in part stimulated by Aat_β_ in an unspecific, ligand-independent manner.

## Discussion

Here we investigated *A. tumefaciens* Aat autotransporter as a candidate scaffold for the display of synthetic passenger domains, aiming at designing a custom, target-specific adhesion system. We repurposed it by fusing the scaffold to lectins, RGD and a VHH. VHH are extremely versatile and can be engineered towards almost any biomarker of interest^56^. As a proof of concept, we rewired *A. tumefaciens* using a cysteine-free VHH anti-GFP to target soluble and cell-displayed GFP.

In animal cells, only low *Agrobacterium*-mediated T-DNA transfection efficiencies have been reported^57,58^. In recalcitrant plants, several reports demonstrate a positive correlation between adhesion and T-DNA delivery efficiency^11,59^. Hence, one possible explanation of the limited progress in *Agrobacterium*-mediated delivery in mammalian cells might be the low affinity of bacteria to such non-natural hosts. We showed that our novel Aat_β_ scaffold has a strong potential in engineering *A. tumefaciens* adhesion to non-natural hosts, including mammalian cells. Thus, our design opens the possibility to further examine the possibility of *A. tumefaciens*-mediated DNA transfer to non-plant hosts. In addition, it provides synthetic biologists with a novel tool to program adhesion of complex multispecies microbial consortia, or the display of enzymes at the surface of other alphaproteobacteria that could be used as additives features in biotechnology or bioremediation^14^.

We next developed a highly sensitive split luciferase assay that enabled us to monitor VirE2 delivery into mammalian cells. The VirE2 transfer in a cell-cell contact-dependent but partly GFP-independent manner has yet to be resolved. Bacteria displaying VHH might release VirE2 upon contact, potentially by VHH display-facilitated lysis. Like in plants, VirE2 could then trigger clathrin-mediated endocytosis^60^. Our results however provide a strong basis for further engineering adhesion to improve *A. tumefaciens* DNA transfer to non-natural host.

Improved *A. tumefaciens-*mediated protein delivery is of importance, as scientists repurposed the T4SS of the bacterium for protein delivery and gene editing, as an alternative to using T-DNA. As an example, Vergunst *et al*. fused VirE2 to the Cre recombinase and leveraged minute amount of proteins transferred to the target cells. They monitored Cre-mediated recombination in plant cells by conferring resistance to cells undergoing recombination^61^. More recently, Cas9 fusions to the VirF peptide responsible for translocation allowed Schmitz *et al*. to target both yeast and plant reporter cells expressing gRNA^62^. Consequently, future experiments could consist in trying such fusions in combination with the adhesin display system to optimize helper proteins delivery at high throughput.

*A. tumefaciens* is an attractive candidate for gene delivery to human cells. The availability of safe, efficient and precisely targeting gene delivery vectors is currently one of the main limiting factors for gene editing therapies *in vivo*. Adeno-associated viruses (AAVs) are the current gold standard for gene delivery. Due to their limited packaging capacity, AAVs however remain a bottleneck for the development of therapies based on CRISPR/Cas9^63,64^. On the other hand, wild-type *A. tumefaciens* injects T-DNA of 25 kb, a size that can easily accommodate several times the CRISPR machinery and repair templates. We thus anticipate that, granted important engineering efforts, *A. tumefaciens* will constitute a powerful tool as an alternative therapeutic DNA delivery vector into human cells.

## Material and methods

Chemicals are purchased from Sigma, unless otherwise stated.

### Cloning

In silico cloning was performed using Benchling software and the cloning strategy of individual plasmids is described in Supplementary table 2.

Phusion polymerase (Thermo) was used for PCR using primers from Microsynth (Switzerland) and restriction enzymes (NEB) for digestion. Plasmid, genomic and gel-purified DNA was extracted using commercially available kits. We performed Gibson assembly using NEB HiFi DNA assembly kit, or classical ligation using the T4 Ligase (Thermo). We noted an improvement of the Gibson assembly efficiency when gel-purifying DNA using the Monarch kit (NEB) compared to other kits. Constructs were transformed by heat shock in XL10Gold (Agilent) prepared in 100 mM calcium chloride and 15% glycerol. Bacteria were plated on Luria broth (LB) plates containing the respective antibiotics and plasmids were screened by Sanger sequencing (Microsynth).

### Composition of home-made media and agar plates

- 20x AB salts (per 200mL): 4 g NH4Cl, 1.2 g MgSO_4_·7H2O, 0.6 g KCl, 0.04 g CaCl_2_, 10 mg FeSO_4_·7H_2_O. Sterile-filtered.
- 20x AB buffer (per 200mL): 12 g K_2_HPO_4_, 4 g NaH_2_PO_4_, pH to 7.0 using either KOH or H_3_PO_4_, as required, before autoclaving.
- Induction medium (IM): 1x AB salts, 0.5% glucose, 2 mM phosphate buffer pH 5.6, 50 mM 2-(4-morpholino)-ethane sulfonic acid (MES)
- Agrobacterium minimal medium: 1x AB salts, 1x AB buffer, 0.5% sucrose, antibiotics.
- ATGN plates: 1x AB salts, 1x AB buffer, 1% glucose, 1.5% noble agar, antibiotics
- ATSN plates: 1x AB salts, 1x AB buffer, 5% sucrose, 1.5% noble agar.
- SDCAA: 18.2% Sorbitol, 2% Glucose, 0.67% Yeast Nitrogen Base, 0.5% Casamino Acids, 0.54% Disodium Phosphate, 0.86% Monosodium Phosphate (Add 1.5% Agar for plates.
- SGCAA: 18.2% Sorbitol, 0.8% Glucose, 8% galactose, 0.67% Yeast Nitrogen Base, 0.5% Casamino Acids, 0.54% Disodium Phosphate, 0.86% Monosodium Phosphate

### Bacterial culture and induction

*E. coli* were cultured at 37°C in LB containing either 100 µg/mL ampicillin (Huberlab), 50 µg/mL kanamycin, 25 µg/mL chloramphenicol, 50 µg/mL spectinomycin (Chemie Brunschwig).

In this study, we used the disarmed strain *A. tumefaciens* C58C1 pMP90 (GV3101, see Supplementary table 3, genome accession number: GCA_000092025.1^65^). *A. tumefaciens* were cultured at 28-30°C in LB containing 60 µg/mL gentamycin (Biochemica) or, when required, either 50 µg/mL kanamycin, 50 µg/mL spectinomycin or 100 µg/mL carbenicillin.

Unless otherwise stated, *A. tumefaciens* was inoculated at an optical density of 0.1 for 8h in LB containing antibiotics and early stationary cells were induced overnight by addition of one volume of induction medium IM and 100 µM acetosyringone (AS) for virulence induction using the VirE promoter. For cumic acid-inducible constructs, early stationary cells were induced overnight in LB by addition of cumic acid at 10 µM.

### Bacterial strain engineering

For replicative plasmids, 1 mL of exponential culture of bacteria was washed 3 times in bi-distilled water, concentrated 20 times and electroporated with 100 ng of plasmid in 1 mm cuvette, rescued for 60 min in Super Optimal broth with Catabolite repression (SOC) medium and plated on the corresponding antibiotics plates. Electro-competent bacteria were snap-frozen in 15% glycerol solution.

For markerless genetic engineering of *A. tumefaciens*, we followed Morton and Fuqua, 2012^66^, using *E. coli* S17-1 for conjugation and pNPTS138 suicide vector (see supplementary Tables 2 and 3), with the following modifications: we added rifampicin (Axon Lab) at 25 µg/mL during selection and counterselection steps (ATGN and ATSN plates) to better kill donor *E. coli*. As kanamycin is inhibited by phosphate-buffered media, we increased the concentration to 1200 µg/mL during selection. LB plates containing rifampicin at 25 µg/mL and kanamycin at 300 µg/mL were sometimes more efficient than the aforementioned ATGN plates. Mutants were screened by colony-PCR using primers flanking the knockin or knockout sites and validated by Sanger sequencing.

### Bacterial staining, titration and quantification

Bacteria displaying VHH were washed with PBS and stained with recombinant GFP at 100 µg/mL for 10 minutes prior to two PBS washes. Bacteria harboring an HA tag were washed with PBS and stained with anti-HA antibody conjugated with FITC (Abcam ab1208) at 10 µg/mL for 75 minutes in the dark on ice, washed once with PBS. For mScarlet-negative cells, viability was checked by concomitant addition of 10 µg/mL propidium iodide (PI) during staining. Wide field fluorescent pictures of bacteria on a coverslip under 1% agarose PBS pads were taken at 100x and 1.5x lens magnification.

### Bioinformatics and modeling

Protein sequences were submitted to the deep learning structure prediction online server RoseTTaFold, provided by the Baker lab: robetta.bakerlab.org ^28^.

Protein sequences were submitted to the online deep neural network software SignalP-5.0 for signal peptide prediction (<u>services.healthtech.dtu.dk/service.php?SignalP-5.0</u>)^29^. Average amino acid usage was extracted from <u>kazusa.or.jp/codon/cgi-bin/showcodon.cgi?species=260551</u>.

### Engineering of S. cerevisiae

For GFP display, we fused eGFP to Aga2p under the control of a galactose-inducible promoter pGal1. Aga2p form disulfide bonds with Aga1p anchored in the cell wall, resulting in eGFP being anchored in the cell wall.

*S. cerevisiae* eby100 was retransformed using 1 µg of pGal1 – Aga2p_eGFP following the protocol of the EZ yeast transformation kit II (Zymo). 100 µL of cells were selected on SDCAA plates, which do not contain tryptophan. Colonies were directly selected by induction and visualization with fluorescent microscopy.

### A. tumefaciens binding to S. cerevisiae

Yeast GFP display induced overnight in SGCAA were washed twice in PBS to remove GFP in suspension and concentrated 10 times. Induced *A. tumefaciens* were washed once in PBS and added to concentrated yeast at a 1 to 1 volume ratio for 60-90 min. Five µL of the cell mixture were transferred to and sandwiched between two coverslips, the yeast cells were left to settle down and imaging was performed at 100x with a confocal microscope and 0.3 µm step. We used NIS Elements (Nikon) for three-dimensional rendering of z-stack pictures and manual cell counting. Per biological replicate, a total of 20 to 50 yeast cells were considered on four different fields of view. For each field of view, the number of bound A. tumefaciens was divided by the number of yeast cell considered. The mean of these values was accounted for one biological replicate.

### Mammalian cell culture and engineering

HEK293T and HeLa cells were cultured in high-glucose Dulbecco’s Modified Eagle Medium (DMEM, Thermofisher) supplemented with 10% FBS (Life Technologies) at 37°C and 5% CO_2_. A GFP-displaying monoclonal HEK cell line was generated by transfecting PvuI-linearized pXP145 and FACS sorting after 7 days of culture.

HeLa cells stably expressing the doxycycline-inducible GFP display were further engineered to stably express LgBit by transduction (pXP499), using lentivector as described previously^47^. After removal of the lentiviruses, a polyclonal cell line was obtained by selection with puromycin (Labforce) at 2 µg/mL for one week.

### Mammalian cell transient transfection

Mammalian cells were transfected with Lipofectamine 3000 (Life Technologies) overnight with 100 ng of purified plasmid per well of 96-well plates, following the manufacturer’s instructions.

### A. tumefaciens binding to mammalian cells

Mammalian cells were washed with medium twice prior to the addition of 10 µL of bacteria induced with 100 µM AS per well of 96-well plates. Bacteria were homogenized by pipetting and left to adhere for 5 hours at 30°C and 5% CO2. For the prevention of binding using soluble GFP, bacteria were incubated with 100 µg/mL recombinant eGFP for 5 minutes prior to the addition to mammalian cells. Consequently, recombinant eGFP was also present during coculture at a concentration of 10 µg/mL. After coculture, wells were washed 5 times with mammalian culture medium and imaging was performed in the region of the well the closest to the dispensing of the medium. Biological replicates were performed on different days and one biological replicate is as follows: confocal microscope Z-stacks were acquired over 12 µm and 2 µm steps in three to five representative fields of view. On each field of view, we estimated the number of HeLa cells and counted bacteria on maximum intensity projection using ImageJ and Trackmate^67^.

### Split luciferase assays

A polyclonal cell line of HeLa cells stably expressing LgBit and doxycycline-inducible GFP display was used. Cells were seeded in 96-well plates (Costar 3603). 6 h later, they were induced when required with doxycycline at 1 µg/mL for overnight expression of GFP-display. Cells were washed twice with medium to remove shed GFP from the supernatant and 22 µL of induced *A. tumefaciens* (LB-IM AS 100 µM) were added to 100 µL of culture medium for 2 h. To reduce unspecific binding and VirE2 transfer, wells were washed 3 times with DMEM with 20% LB-IM and 100 µM AS, and infection was prolongated for 3 more hours before readout (Fig. 5D and S8A-B), else infection was performed for 5 hours.

Wells were washed with OptiMEM (Thermo) twice and 20 µL of a 1:19 mixture of substrate:buffer from the Nano-Glo Live Cell Assay (Promega) were added to 80 µL of OptiMEM per well. After 3 min incubation with the reagent, luminescence activity was acquired using a multiwell plate reader (Tecan Spark) with 5 s integration time per well. Background from wells with only reagent and OptiMEM was subtracted from the values.

Biological replicates were performed on different days and one biological replicate consists in three technical replicates (separate wells).

### Microscopy

For widefield visualizations of bacteria, we used a Nikon TiE epifluorescence microscope equipped with a Hamamatsu ORCA Flash 4 camera and an oil immersion 100x Plan APO N.A. 1.45 objective.

For bacterial adhesion to yeast and mammalian cells, we used a Nikon Eclipse Ti2-E inverted microscope coupled with a Yokogawa CSU W2 confocal spinning disk unit and equipped with a Prime 95B sCMOS camera (Photometrics). We used either a 40x air objective with N.A. of 1.15 to acquire z-stacks for mammalian cells or a 100x oil immersion objective with N.A. of 1.45 for yeast cells.

### Production of recombinant proteins

6x-His tagged eGFP on a pET28a vector (see Supplementary table 2) was retransformed into BL21 strain. We induced production with 1 mM IPTG (Fisher bioreagents) at room temperature overnight. We pelleted and lysed bacteria by sonication in lysis buffer (Tris 100mM, NaCl 0.5M, glycerol 5%) and eGFP was purified using fast flow His-affinity columns (GE Healthcare) and eluted with 0.5 M imidazole. We exchanged buffer to PBS using 30kDa ultracentrigation spin columns (Merck) and adjusted the concentration to 1 mg/mL. Aliquots were snap frozen for further use.

## Supporting information

Supplementary figure 6: Analysis of the signal peptides of the two putative autotransporters of *A. tumefaciens*

Supplementary figure 7: Removal of the disulfide bonds of the native passenger domain in Atu5364 promotes even surface distribution of the passenger domain.

Supplementary figure 8: VHH display strategy using Atu5364 as scaffold.

Supplementary figure 9: Successful inducible RGD and lectin display at the surface of *A. tumefaciens*

Supplementary figure 10: Schematic of the VHH_GFP__Aat_β_ introduction into the VirE2 locus.

Supplementary figure 11: Synthetic adhesion of *A. tumefaciens* to HEK cells.

Supplementary figure 12: HiBit::VirE2 complements LgBit in reporter cells.

Supplementary figure 13: VHH display causes adhesion-independent VirE2 uptake, which might result from bacterial lysis upon contact.

Supplementary methods and references

## Acknowledgments

The authors are grateful to Dr. Ingmar Riedel-Kruse (The University of Arizona) and Dr. Luis Angel Fernández (Centro Nacional de Biotecnologia) for the nanobody anti-GFP construct, to Dr. Bruno Correia, Stéphane Rosset and Dr. Leo Scheller (Ecole Polytechnique Fédérale de Lausanne) for the production and purification of recombinant proteins, *S. cerevisiae* eby100 strain, NanoLuc and split NanoLuc constructs, to Dr. Csaba Koncz (Max Planck Institute for plant breeding research) for *A. tumefaciens* GV3101.

## Author contributions

X.P. and A. Persat conceived the work. X.P. designed and performed the experiments. A. Pham and J.W. performed experiments for the revision of the manuscript. Z.A-M. provided support for cell culture, DNA purification and colony PCR. The EPFL Flow Cytometry Core Facility performed single cell sorting. X.P. and A. Pham analyzed the data. X.P. and A. Persat wrote the manuscript. All authors gave input and approved the manuscript.

