## Supplementary figures 1-8 for "Engineering *Agrobacterium tumefaciens* adhesion to target cells"

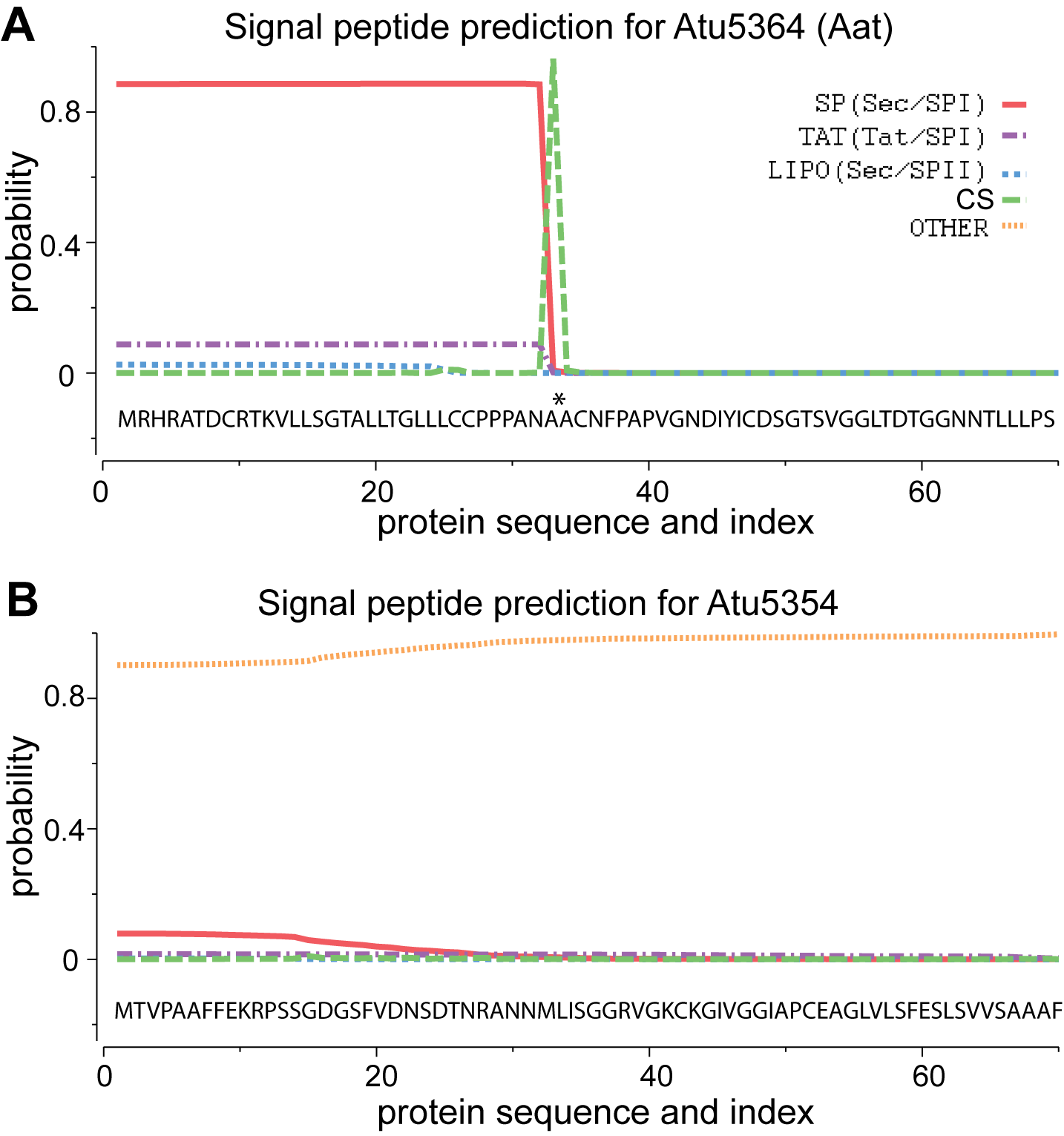

**Supplementary figure 1: Analysis of the signal peptides of the two putative autotransporters of *A. tumefaciens* using SignalP-5.0**^1^**.** (A) Atu5364 (Aat) is the only protein for which a signal peptide is predicted, of a length of 33 amino acids. (B) Atu5354 does not have any predicted signal peptide. Signal peptide prediction using SignalP - 5.0. Sec/SPI: "standard" secretory signal peptides transported by the Sec translocon and cleaved by Signal Peptidase I (Lep). Sec/SPII: lipoprotein signal peptides transported by the Sec translocon and cleaved by Signal Peptidase II (Lsp). Tat/SPI: Tat signal peptides transported by the Tat translocon and cleaved by Signal Peptidase I (Lep). CS: putative cut site.

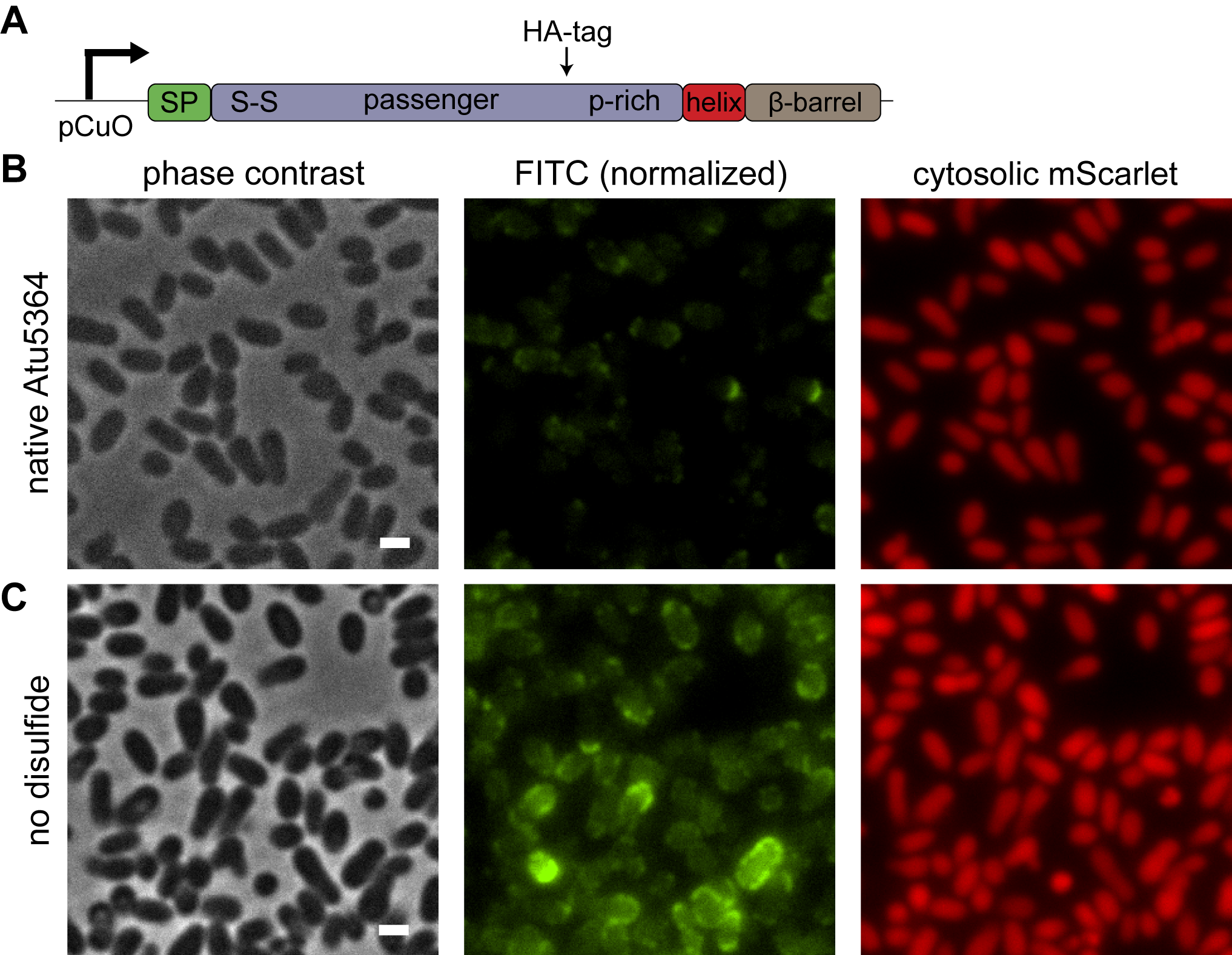

**Supplementary figure 2:** **Removal of the disulfide bonds of the native passenger domain in Atu5364 promotes even surface distribution of the passenger domain.** (A) Schematic of the genetic constructs used for assessing the impact of Atu5364’s disulfide bond on display. pCuO: synthetic cumic acid-inducible promoter. S-S: native disulfide bond. P-rich: proline-rich linker. (B-C) *A. tumefaciens mScarlet atu5364-* was retransformed with a cumic acid-inducible *atu5364* internally tagged with HA (pCuO - atu5364_HA) (B) or pCuO - atu5364_cysteine_free_HA (C) and stained with FITC-conjugated anti-HA antibody (anti-HA-FITC). (B) The staining is weak or unipolar. (C) Disulfide-free Atu5364 is strongly expressed at the surface of bacteria. Intensity scale is identical between samples. Bars, 1 µm.

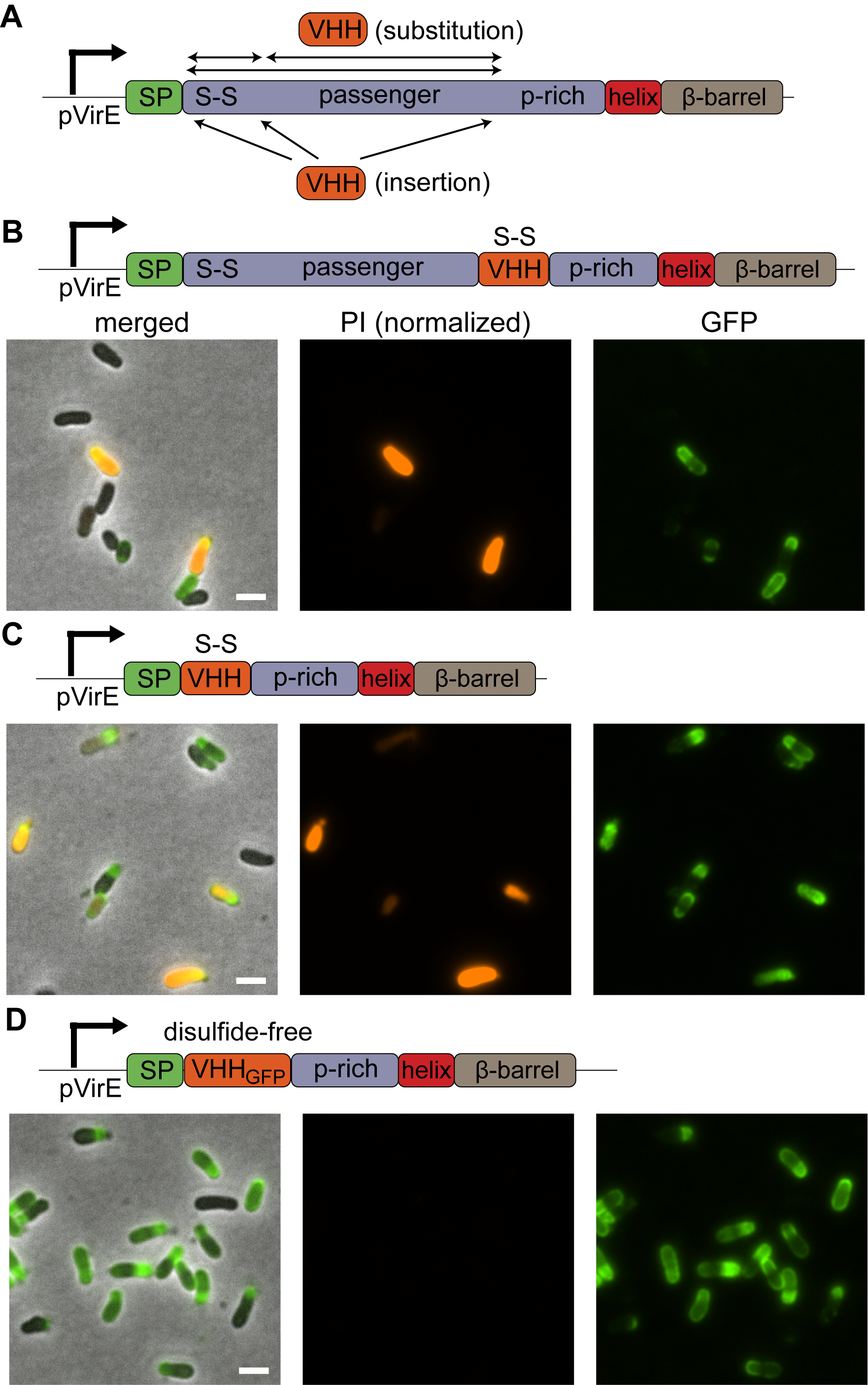

**Supplementary figure 3: VHH display strategy using Atu5364 as scaffold.** (A) Schematic depicting the general VHH anti-GFP introduction approach: VHH either replaces the disulfide-bond (S-S)-containing part of the passenger domain, the remaining part or the whole passenger domain, proline-rich (p-rich) linker excluded (amino acids 35-160, 161-512 or 35-512). VHH is also introduced after the signal peptide (SP), after the S-S of before the p-rich linker. (in front of C35, D161 or A513). (B, C) *A. tumefaciens* does not efficiently display VHH harboring its native disulfide bond. *A. tumefaciens* was retransformed with pVirE - internalVHH display (B), pVirE – VHH display (C). (D) Removal of the disulfide bond by mutagenesis of VHH C24A and C98V (VHH_GFP_) increases display efficiency and retains affinity to GFP. Dead cells were stained with propidium iodide (PI) and VHH was stained with recombinant GFP. PI intensity scale is identical between samples. Bars, 2 µm.

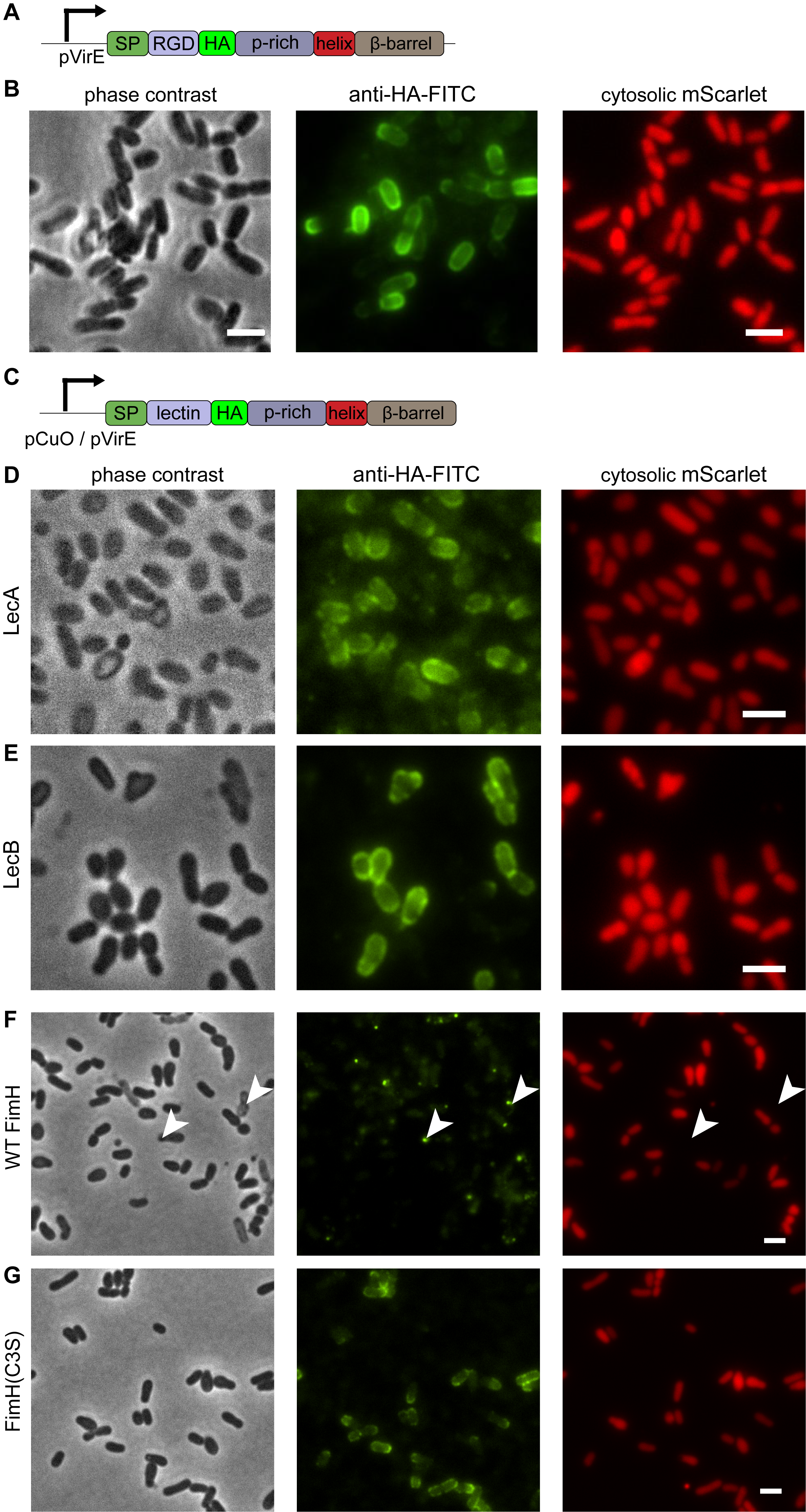

**Supplementary figure 4: Successful inducible RGD and lectin display at the surface of *A. tumefaciens.*** (A) Schematic of the arginine-glycine-aspartic acid (RGD) fused to HA tag (HA) display introduction under a VirE promoter (pVirE). (B) *A. tumefaciens* mScarlet was retransformed with pVirE – RGD_HA display and induced with acetosyringone for 8h prior to FITC-conjugated anti-HA staining (FITC). Bars, 1 µm. (C) Schematic of the display approach: lectins fused to an HA tag are replacing the passenger domain in Atu5364 under a cumic acid-inducible promoter (pCuO) or pVirE. (D-E) Lectins from *P. aeruginosa* LecA and LecB are placed under pCuO. *A. tumefaciens* mScarlet was retransformed with pCuO – LecA_HA display (D) or pCuO – LecB_HA display (E), induced and successfully stained with FITC-conjugated anti-HA antibody (anti-HA-FITC). (F-G) FimH j96 from uropathogenic *E. coli* was placed under pVirE. (F) Uropathogenic *E. coli* FimH (strain j96) display induces cell death and patchy HA-staining. *A. tumefaciens* mScarlet was retransformed with pVirE – FimH_HA tag display, induced and stained with anti-HA-FITC. Low cell viability is visible in phase contrast (light grey cell bodies) and mScarlet (no signal). The anti-HA staining is uneven and mostly correlates with dead bacteria (white arrowheads). (G) FimH(C3S) display increases homogeneity of HA-staining and viability. *A. tumefaciens* mScarlet was retransformed with pVirE – FimH(C3S)_HA tag display, induced and stained with FITC-conjugated anti-HA antibody (anti-HA-FITC). Bacteria are healthy and the anti-HA staining is evenly distributed around the outer membrane. Bars, 2 µm.

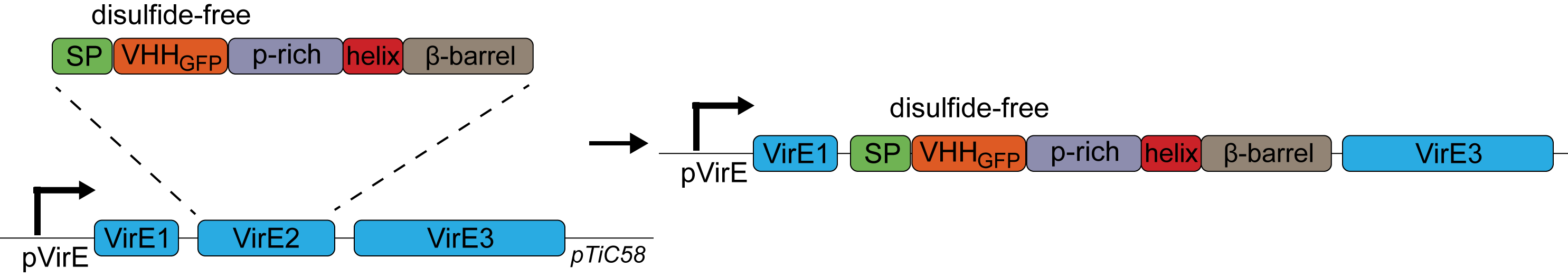

**Supplementary figure 5: Schematic of the VHH_GFP__Aat_β_ introduction into the VirE2 locus.** Using a two-step allelic exchange protocol, VirE2 was replaced by the VHH_GFP__Aat_β_ construct on the pTiC58 megaplasmid.

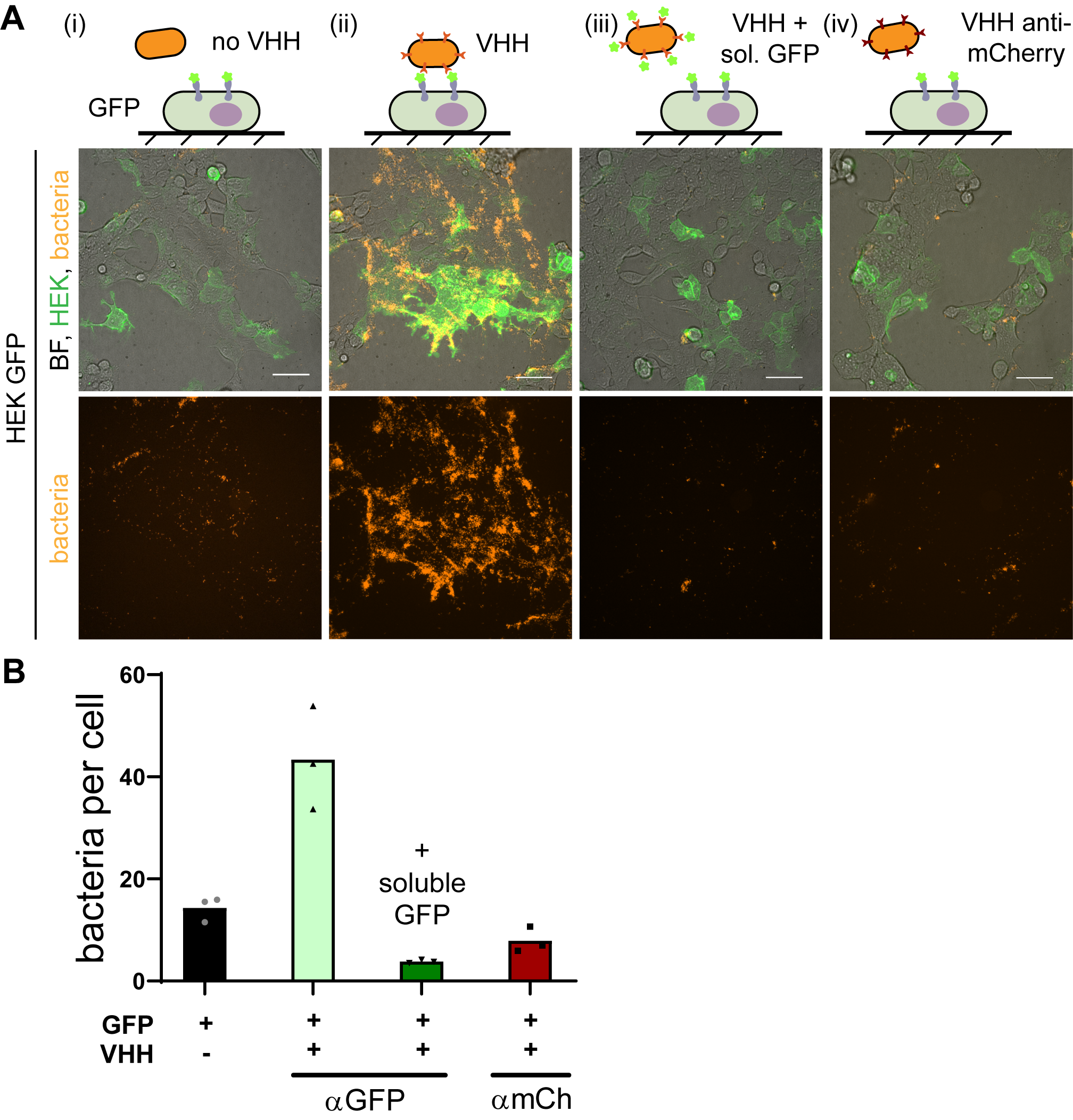

**Supplementary figure 6: Synthetic adhesion of *A. tumefaciens* to HEK cells.** (A) Representative maximum intensity projections of *A. tumefaciens* (orange) binding to GFP-displaying HEK cells (green) in the different conditions. (i) no VHH_GFP__Aat_β_ and GFP-display, (ii) VHH_GFP__Aat_β_ and GFP display (iii) VHH_GFP__Aat_β_ and GFP-display prevented by soluble GFP and (iv) VHH_mCherry__Aat_β_ and GFP display BF, bright field (greyscale). Bars, 50 µm. (B) Quantification of the average number of bacteria per GFP-displaying HEK cells. *A. tumefaciens* was retransformed with an empty vector (VHH -), pVirE – VHH_mCherry_ (αmCh) or pVirE – VHH_GFP_ (αGFP). In the last column, soluble recombinant eGFP was added to prevent binding by saturating VHH receptors. Bars represent the mean of technical triplicates.

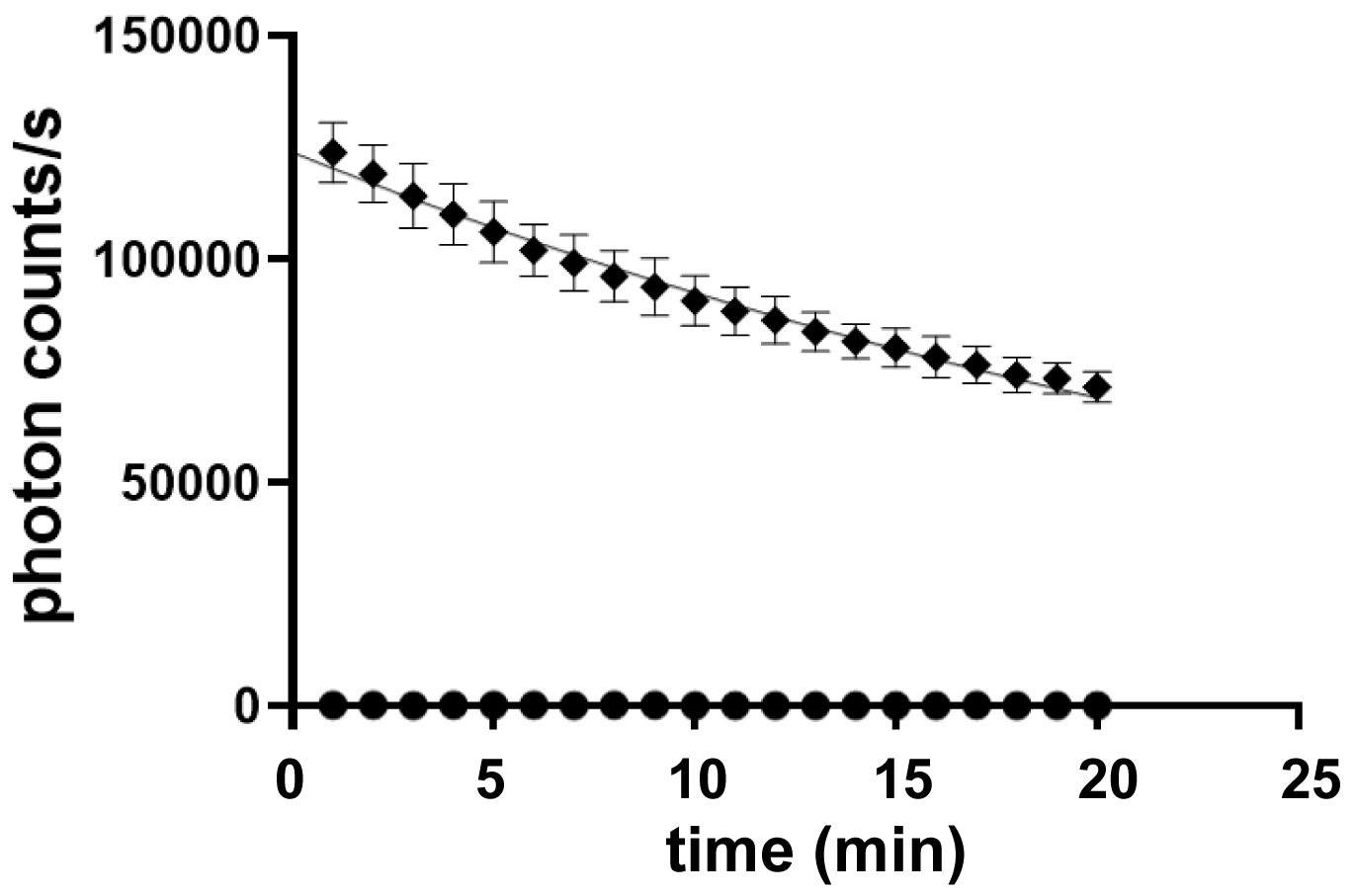

**Supplementary figure 7: HiBit::VirE2 complements LgBit in reporter cells.** GFP-displaying HeLa cell line stably and constitutively expressing LgBit were transiently transfected with a plasmid encoding pCMV - HiBit::VirE2 expression (diamonds) or empty vector (circles). Live cell luminescence was measured over time and an exponential decay function was fitted to the data, highlighting a half-life of the signal of 24 min. This half-life is more than 4 times lower than reported by the manufacturer, and might be due to substrate depletion caused by a high concentration of reconstituted enzyme. Error bars represent the standard deviation of technical triplicates.

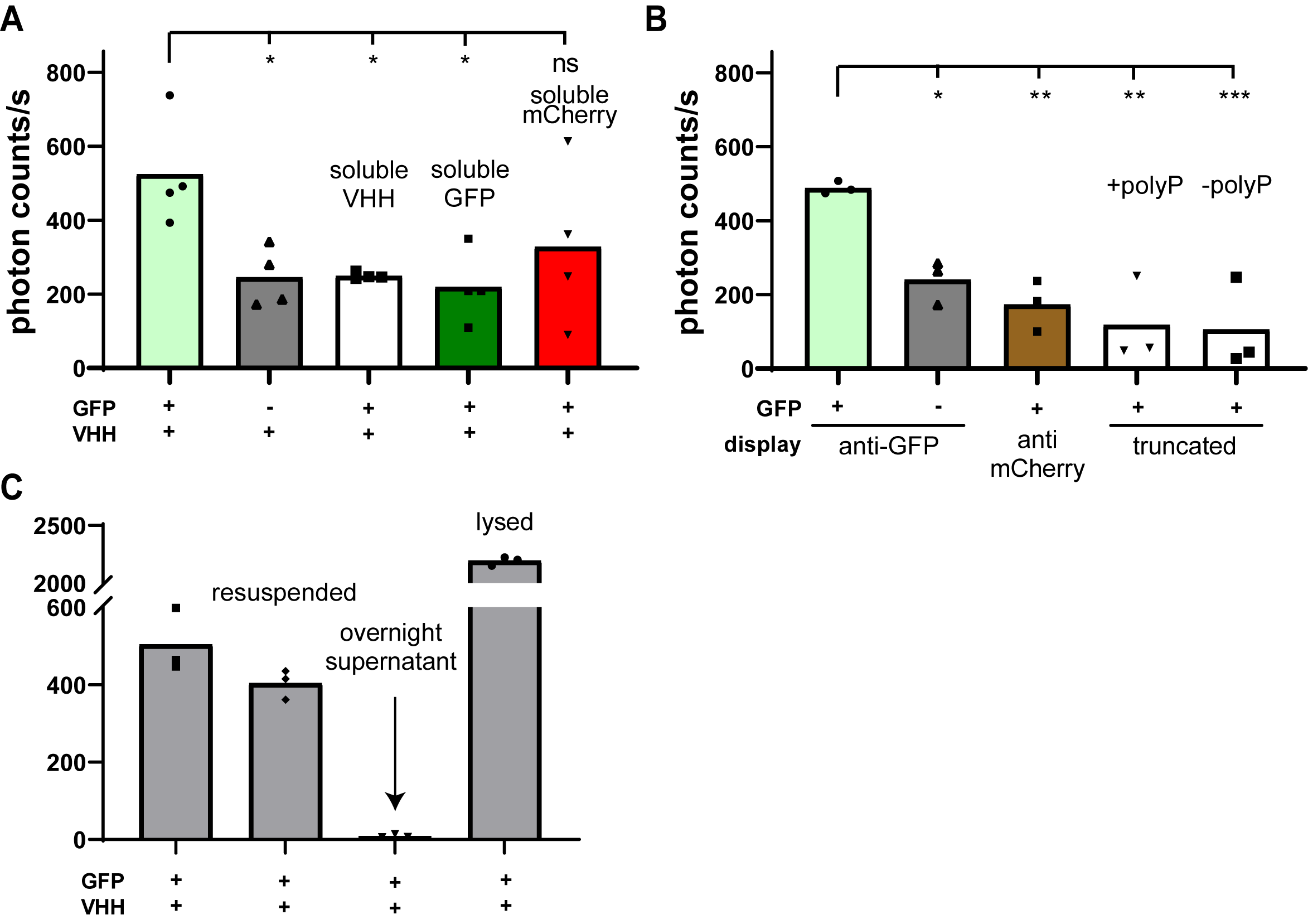

**Supplementary figure 8: VHH display causes adhesion-independent VirE2 uptake, which might result from bacterial lysis upon contact.** (A-B) The display of VHH has an intrinsic contribution to VirE2 transfer. Luminescence signal from reporter HeLa cells expressing LgBit and display GFP or not (+ or -) co-cultured with bacteria. (A) *A. tumefaciens* *VHH_GFP_::ΔVirE2* pVirE - HiBit::VirE2 was used in all conditions. Addition of soluble VHH (white column), soluble GFP (dark green) does not abolish luminescence signal. The addition of soluble mCherry (dark red) does not significantly reduce the signal (control). (B) *A. tumefaciens* *VHH_GFP_::ΔVirE2* pVirE - HiBit::VirE2 was used for the first two columns. *A. tumefaciens* *ΔVirE2* pVirE - HiBit::VirE2 was retransformed with pVirE - VHH_mCherry__Aat_β_ Spec^R^ for the third column, with pVirE – linker_Aat_β_ for the fourth and pVirE - Aat_β_ for the fifth column. Statistical tests: one-way ANOVA followed by Dunnett’s post hoc test (*** P<0.001, ** P<0.01, * P<0.05, ns P>0.05). (C) Luminescence signal comes from reporter cells infected by bacteria resuspended in induction medium and not their overnight supernatant, however, freshly lysed bacterial supernatant increases the signal. “resuspended”: *A. tumefaciens* *VHH::ΔVirE2* pVirE - HiBit::VirE2 were pelleted and resuspended in fresh induction medium, added to the GFP-displaying HeLa cells expressing LgBit. “overnight supernatant”: the supernatant of the aforementioned pelleted bacteria is added to mammalian cells. “Lysed”: Bacteria were lysed by sonication on ice, spun down and the supernatant was applied to mammalian cells. Datapoints represent the means of biological triplicates.

**Supplementary methods**

*Lysis of A. tumefaciens* by sonication

Induced cultures were sonicated in 1.5 mL Eppendorf on ice using a Branson 550 sonicator equipped with a microprobe at 30% power. 3 seconds pulse and 10 seconds rest cycles were applied for a total time of 45 seconds of sonication.

*Production of recombinant proteins*

6x-His tagged mCherry on a pET28a vector (see Supplementary table 2) was retransformed into BL21 strain. We induced production with 1 mM IPTG (Fisher bioreagents) at room temperature overnight. We pelleted and lysed bacteria by sonication in lysis buffer (Tris 100mM, NaCl 0.5M, glycerol 5%) and eGFP was purified using fast flow His-affinity columns (GE Healthcare) and eluted with 0.5 M imidazole. We exchanged buffer to PBS using 30kDa ultracentrigation spin columns (Merck) and adjusted the concentration to 1 mg/mL. Aliquots were snap frozen for further use.

*Recombinant VHH*

Recombinant VHH anti-GFP was purchased from LuBioScience (reference GT-250).
