## Supplementary tables 1-3 for "Engineering *Agrobacterium tumefaciens* adhesion to target cells"

**Table S1: Overview of the top templates used by SwissModel and I-Tasser homology-model modeling software with the passenger domain of Atu5364 (aa 35-501) as input.**

Here, different fragments of 10 to 30% of the query sequence were aligned to templates. The “Local % identity” represents the identity percentage of these fragments with the templates used for modeling.

| Protein template name | Host | Local % Identity | Function |
| --- | --- | --- | --- |
| IcsA/VirG autotransporter | *Shigella flexneri* | 34 | Intracellular, actin-based motility^1^ |
| P.69 pertactin autotransporter | *Bordetella (para)pertussis* | 30 | Binding to host cells^2^ |
| Antifreeze protein | *Marinomonas primoryensis* | 27 | Binding to ice crystal and ice growth prevention^3^ |
| RsaA | *C. crescentus* | 27 | S-layer production for protection and or pathogenicity^4,5^ |
| Ag43 autotransporter | *E. coli* | 24 | Self-association^6^ |

**Supplementary table 2: plasmids used in this study.**

Construction by Gibson assembly unless stated otherwise.

| Internal reference | Plasmid name(s) | Description  Cloning approach:   - - Templates, primers used for PCR and/or enzymes for digestion - Followed by Gibson assembly or T4 ligation when all DNA fragments are digested | Reference |
| --- | --- | --- | --- |
| 290 | pcDNA3.3 eGFP | Mammalian vector for pCMV-driven eGFP expression. Amp^R^, Neo^R^ | Addgene 26822 ^7^ |
| 296 | pFGL815 | Binary vector backbone for *Agrobacterium tumefaciens* with pVS1 ori. Kan^R^ | Addgene 52322 ^8^ |
| 302 | pPZP200 | *A. tumefaciens* binary vector with an MCS within T-DNA. pVS1 ori. Spec^R^ | Hajdukiewicz *et al*., 1994 ^9^ |
| 304 | pDSG339  pSB3K3 TetR pTet Neae2_VHH | Tetracycline-inducible VHH anti-GFP display based on a truncated intimin autotransporter scaffold (*E. coli*). Kan^R^ | Glass et. al, 2018 ^10^ |
| 368 | pCB301  pJL-TRBO-G | We used the backbone of this binary vector in *A. tumefaciens* with IncPa1 ori. Kan^R^ | Addgene 80083 ^11^ |
| 372 | pGEX6P1-mCherry-Nanobody | *E. coli*-optimized GST-tagged Nanobody anti mCherry | Addgene 70696 ^12^ |
| 433 | pGB2-24 FimH j96 | *E. coli* vector for expression of FimH-j96 variant in KB18. Chloramph^R^ | Sokurenko *et al*, 1994 ^13^ |
| 491 | pCTcon2 | Yeast display expression vector with the yeast *trp1* selection marker. Kan^R^  For in-frame insertion of C-terminal fusions to the cell-wall anchoring protein Aga2p, under the control of a galactose-inducible promoter. | Addgene 41843 ^14^ |
| 522 | pET21a_CymR | *E. coli* expression vector for CymR production. Amp^R^ | Addgene 51165 ^15^ |
| 523 | pKD227 | Expresses *S. baltica* TtrS under the Ptac promoter. Used to PCR out the Pkm promoter . Spec^R^ | Addgene 90954 ^16^ |
| 524 | pJM101 | miniTn7 delivery plasmid with lacIq-Ptac inducible promoter. Amp^R^ Genta^R^ | Addgene 110558 ^17^ |
| 1159 | pRRL-PGK-LacZ-3HA-IRES-puro | Lentivector with constitutive hPGK promoter, LacZ HAtag, IRES. Amp^R^, Puro^R^ | Trono lab, EPFL, unpublished |
| pXP135 | pVirE-VirE0-1-2 | VirE0-1-2 gene in pFGL815 (resolvase excluded). Kan^R^   - - pFGL815, agacggcggatacgttgccagccagccaacagct + ccgtaaacagtctgtagaaccgcaccaggacggc - *- A. tumefaciens* gDNA, AGCTGTTGGCTGGCTGGCAACGTATCCGCCGTCT + gccgtcctggtgcggttctacagactgtttacggttg | This study |
| pXP145 | pCMV – GFP display | Constituve eGFP (N105Y, E125V,Y146F) display based on a truncated C-terminal CD80 anchor, Amp^R^, Neo^R^. | Pierrat et. al, 2021 ^18^ |
| pXP155 | pVirE - mScarlet | mScarlet under control of VirE promoter in pFGL815. Kan^R^   - - pXP135, CTCGCCCTTGCTCACCATcgtctaactccttttagccggc + GACGAGCTGTACAAGTGAaaccgcaccaggacggcc - - pZA002 (mScarlet), ATGGTGAGCAAGGGCGAG + TCACTTGTACAGCTCGTCC | This study |
| pXP166 | pVirE - internalVHH display | Native VHH anti-GFP introduced between E512 and A513 into *Aat_β_* under control of VirE promoter in pFGL815. Kan^R^   - - pXP182, cccaggtcaccgtctcctcagctgaggctcccgatat + accagctgcacctgagccatctccggggtggtatttgg - - pDSG339, atggctcaggtgcagct + tgaggagacggtgacct | This study |
| pXP169 | pVirE – VHH display | Native VHH anti-GFP replacing the passenger domain of atu5364 from C35 to E512 included, under control of VirE promoter in pFGL815. Kan^R^   - - pXP182, cccaggtcaccgtctcctcagctgaggctcccgatat + accagctgcacctgagccatagctgcgtttgcgggag - - pDSG339, atggctcaggtgcagct + tgaggagacggtgacct | This study |
| pXP182 | pVirE - atu5364 WT | Atu5364 WT under control of VirE promoter in pCB301. Kan^R^   - - pXP135, aaccgcaccaggacggc + cgtctaactccttttagccggc - *- A. tumefaciens* gDNA, cggctaaaaggagttagacgatgcggcatcgagctac + ctggccgtcctggtgcggttctaccacttgatgttgagaccg | This study |
| pXP213 | pVirE – VHH_GFP__Aat_β_ in pFGL815 (Figure 2) | VHH anti-GFP cysteine-free (C24A, C98V) a.k.a. VHH_GFP_ display on Aat_β_ under control of VirE promoter in pFGL815. VHH replaces the passenger domain of atu5364 from C35 to E512 included. Kan^R^   - - pXP169, gccgtgtattacGTGaatgtcaatgtgggctttga + tccagaggctgcCGCggagagtctcagagaccc - - pXP169, ctgagactctccGCGgcagcctctggattccc + cacattgacattCACgtaatacacggccgtgtc | This study |
| pXP226 | pET28 - eGFP | 6xHis-tagged eGFP in pET28a for recombinant expression. Kan^R^ | Pierrat et. al, 2021 ^18^ |
| pXP228 | pVirE – VirE2 WT | Plasmid complementation of VirE2 under control of VirE promoter in pCB301. Kan^R^   - - pCB301, caaccgtaaacagtctgtagctaagagaaaagagcgtttattaga + agacggcggatacgttgccagccagccaacagct - - pXP135, caacgtatccgccgtc + ctacagactgtttacggttgg | This study |
| pXP229 | pVirE – VHH_GFP__Aat_β_ in pCB301  (used in Figures other than Figure 2 for ori compatibility in bacteria transformed with two plasmids) | VHH anti-GFP cysteine-free (C24A, C98V) a.k.a. VHH_GFP_ display on Aat_β_ under control of VirE promoter in pCB301. VHH replaces the passenger domain of atu5364 from C35 to E512 included. Kan^R^   - - pCB301, gtctcaacatcaagtggtagctaagagaaaagagcgtttattaga + agacggcggatacgttgccagccagccaacagct - - pXP213, caacgtatccgccgtc + ctaccacttgatgttgagacc | This study |
| pXP233_  pNPTS138 | *virE2* knockout | Suicide vector for the knockout of *virE2* in *A. tumefaciens*. pNPTS138, two-step allelic exchange protocol. Kan^R^ Sucrose^S^   - - pNPTS138, EcoRI + XbaI - Overlap extension PCR of the following two PCR products followed by EcoRI + XbaI digestion:   - - pXP229, atgaccatgattacgaattcagtcaagttgacctggcg + catcgtctaactccttttagcc   - - *A. tumefaciens* gDNA, ctaaaaggagttagacgatgtagatttcctgataccgcgtc + gcctgcaggtcgactctagatatttcacatgcttcgcgc | This study |
| pXP234_  pNPTS138 | *VHH_GFP__Aat_β_* knockin instead of *virE2* | Suicide vector for the knockin of the gene coding for cysteine-free VHH anti-GFP display on Aat_β,_ instead of *virE2* in *A. tumefaciens*. pNPTS138, two-step allelic exchange protocol. Kan^R^ Sucrose^S^   - - pNPTS138, EcoRI + XbaI - Overlap extension PCR of the following two PCR products followed by EcoRI + XbaI digestion:   - - pXP229, atgaccatgattacgaattcagtcaagttgacctggcg + ctaccacttgatgttgagacc   - - *A. tumefaciens* gDNA, gtctcaacatcaagtggtagatttcctgataccgcgtcag + gcctgcaggtcgactctagatatttcacatgcttcgcgc | This study |
| pXP250 | pSynth - mScarlet knock-in in *tetR* | Suicide vector for the introduction of *mScarlet* under control of a consensus synthetic promoter originally optimized for *E. coli* : <http://parts.igem.org/Part:BBa_J23119> , introduced into the *tetR* locus. pNPTS138, two-step allelic exchange protocol.Kan^R^ Sucrose^S^   - - pNPTS138, EcoRI + HindIII - *- A. tumefaciens* gDNA, cggccgaagctagcgaattcccttcatttcggctgttcac + AGGACTGAGCTAGCTGTCAAcggtatgaaggagaagctgc - - pZA002, TTGACAGCTAGCTCAGTCC + TTACTTATACAGTTCATCCATACCAC - *- A. tumefaciens* gDNA, TGGATGAACTGTATAAGTAAgcactttatcttctcgccttg + agccggctggcgccaagcttggtattggcggcggtattatc | This study |
| pXP255 | pVirE – VHH_mCherry__Aat_β_ | VHH anti-mCherry cysteine-free (C24A, C98V) display on Aat_β_ under control of VirE promoter in pCB301   - - pXP229, tatattagcagcaatcagcgcctgtatggttactggggccagggg + gctttctgcaaaacgtccagaggctgcCGC - - pGEX6P1-mCherry-Nanobody, GCGgcagcctctggacgttttgcagaaagcagc + ctgattgctgctaatatagttacccagattggctgcCACataatacac | This study |
| pXP256 | pET28 - His6_mCherry | 6xHis-tagged mCherry in pET28a for recombinant expression. Kan^R^  - pET28a* ATGGACGAGCTGTACAAGTGAGATCCGGCTGCT + CTCGCCCTTGCTCACCATGCTGCTGTGATGATGATG  - mCherry-Tubulin-C-18** ATGGTGAGCAAGGGCGAG + TCACTTGTACAGCTCGTCC | *EMD Biosciences  **Addgene 55148 |
| pXP267 | pGal1 – Aga2p_eGFP | Galactose-inducible Aga2p-eGFP yeast display in pCTcon2, Kan^R^, *trp1*  Ligation of the following:   - - pCTcon2, NheI + XhoI - - pcDNA3.3 egfp, tcggctagcatggtgagcaagggcgag + gatctcgagtcacttgtacagctcgtcc followed by NheI + XhoI | This study |
| pXP269 | pVirE – RGD display (no HA) | RGD display on Aat_β_ under control of VirE promoter in pCB301. AG*RGD*SP replaces the passenger domain of atu5364 from C35 to E512 included. Kan^R^   - - pXP182, CGCGGCGACAGCCCGgctgaggctcccgatatcac + GCCGGCagctgcgtttgcggg | This study |
| pXP306 | pVirE – FimH_HA tag display | FimH(j96)_HA-tag display on Aat_β_ under control of VirE promoter in pCB301. Kan^R^   - - pXP229, TACCCGTATGATGTTCCCGACTATGCCgctgaggctcccgatatc + agctgcgtttgcggg - - pGB2-24 FimH j96, ctcctcccgcaaacgcagctttcgcctgtaaaaccgcc + AACATCATACGGGTAgccgccagtaggcac | This study |
| pXP307 | pVirE – FimH(C3S)_HA tag display | FimH(C3S)(j96)_HA-tag display on Aat_β_ under control of VirE promoter in pCB301. Kan^R^   - - pXP229, TACCCGTATGATGTTCCCGACTATGCCgctgaggctcccgatatc + agctgcgtttgcggg - - pGB2-24 FimH j96, ctcctcccgcaaacgcagctttcgccAGCaaaaccgcc + AACATCATACGGGTAgccgccagtaggcac | This study |
| pXP313 | pCuO - mScarlet | Cumic-acid inducible mScarlet in pFGL815. Tac promoter followed by cumic acid operators (CuO) bound by CymR. Addition of cumic acid detaches CymR from CuO and restores transcription Kan^R^   - - pXP155, ccagccagccaacagc + cagcacactggcggccgttactagaaataattttgtttaactttaagaaggagatataccATGGTGAGCAAGGGCG - - pET21a_CymR, gggagctgttggctggctggttaacgtttgaattttgcataacgt + atgagcccgaaacgtcg - - pKD227, cgacgtttcgggctcatcttgcgaaacgatcctcatcct + gcatgcatttaaatacgcgtccggaattgccagctg - - pJM101, acgcgtatttaaatgcatgctcgactgcacggtgcac + taacggccgccagtgtgctggaattcataatacaaacagaccagattgtctgtttgttccacacattatacgagccgat | This study, based on Denkovskiene *et al*. 2015^19^ |
| pXP319 | pCuO – LecA_HA display | Cumic-acid inducible PAO1 LecA display on Aat_β_ with HA tag in pFGL815. Kan^R^   - - pXP313, AACCGCACCAGGACGG + gtatatctccttcttaaagttaaacaaaattatttc - - pXP229, actttaagaaggagatatacatgcggcatcgagctac + agctgcgtttgcggg - - PAO1 gDNA, ctcctcccgcaaacgcagctATGGCTTGGAAAGGTGAGG + GAACATCATACGGGTAgccgccGGACTGATCCTTTCCAATATTGAC - - pXP229, TACCCGTATGATGTTCCCGACTATGCCgctgaggctcccgatatc + gccgtcctggtgcggttctaccacttgatgttgagaccg | This study |
| pXP320 | pCuO – LecB_HA display | Cumic-acid inducible PAO1 LecB display on Aat_β_ with HA tag in pFGL815. Kan^R^   - - pXP313, AACCGCACCAGGACGG + gtatatctccttcttaaagttaaacaaaattatttc - - pXP229, actttaagaaggagatatacatgcggcatcgagctac + agctgcgtttgcggg - - PAO1 gDNA, ctcctcccgcaaacgcagctATGGCAACACAAGGAGTGTTC + AACATCATACGGGTAgccgccGCCGAGCGGCCAGT - - pXP229, TACCCGTATGATGTTCCCGACTATGCCgctgaggctcccgatatc + gccgtcctggtgcggttctaccacttgatgttgagaccg | This study |
| pXP323 | pCuO - atu5364_HA | Cumic-acid inducible atu5364 display with HA tag in pFGL815. Kan^R^   - - pXP313, NcoI + BsrGI - - pXP313, tagaaccgcaccaggac + gtatatctccttcttaaagttaaacaaaattatttc - - pXP304, actttaagaaggagatatacatgcggcatcgagctac + ctggccgtcctggtgcggttctaccacttgatgttgagaccg | This study |
| pXP324 | pCuO - atu5364_cysteine_free_HA | Cumic-acid inducible cysteine-free Atu5364 (C35M, C48S) display with HA tag in pFGL815. Kan^R^   - - pXP313, NcoI + BsrGI - - pXP313, tagaaccgcaccaggac + gtatatctccttcttaaagttaaacaaaattatttc - - pXP305, actttaagaaggagatatacatgcggcatcgagctac + ctggccgtcctggtgcggttctaccacttgatgttgagaccg | This study |
| pXP340 | tetON – GFP-display | Mammalian lentivector containing a tetracycline-inducible GFP(N105Y, E125V,Y146F) -display anchored with a truncated CD80 transmembrane domain in pRRLSIN.cPPT.insert.WPRE | Pierrat et. al, 2021 ^18^ |
| pXP479 | pVirE – RGD_HA display | RGD + HA tag display on Aat_β_ under control of VirE promoter in pCB301. AG*RGD*SP replaces the passenger domain of atu5364 from C35 to E512 included. Kan^R^   - - pXP269, tagaaccgcaccaggac + AACATCATACGGGTACGGGCTGTCGCCG - - pXP269, TACCCGTATGATGTTCCCGACTATGCCgctgaggctcccgatatca + gccgtcctggtgcggttctaccacttgatgttgagaccg | This study |
| pXP496 | pVirE - HiBit::VirE2 | HiBit peptide introduced at position P61 (determined by sequence alignment to VirE2(pTiBo542) ^20,21^) in *VirE2* under control of VirE promoter in pCB301. Kan^R^   - - pCB301, ctaagagaaaagagcgtttattagaataatcg + agacggcggatacgttgccagccagccaacagct - - pXP135, caacgtatccgccgtc + TTCTTGAACAGGCGCCAGCCCGAGACcgggcttccgtgcatg - - pXP135, GGCTGGCGCCTGTTCAAGAAGATCTCGactcacacggatgatctcgg + taaacgctcttttctcttagctacagactgtttacggttggg | This study |
| pXP499 | hPGK – LgBit in lentivector | Second generation lentivector containing a constitutive hPGK promoter driving LgBit (split NanoLuc) expression. Amp^R^, Puro^R^  Ligation of the following:   - - pRRL-PGK lacZ-3HA-IRES-puro, BamHI + NheI - - LgBit (synthesized), tctctccccAGGGGGATCCACCatgGTCTTCACACTCGAAGATTTCG + CGGCCGCGTTTCGCTAGCttaGTTGATGGTTACTCGGAACAG, then BamHI + NheI | Dixon et. al, 2016 ^22^. This study. |
| pXP507 | pCMV – HiBit::VirE2 | HiBit peptide introduced at position P61 (determined by sequence alignment to VirE2(pTiBo542) ^20,21^) in *VirE2* under control of a mammalian CMV promoter in. Amp^R^/Neo^R^   - - pcDNA3.3 eGFP, gctgccttctgcgggg + ggtggctcttatatttcttcttactcttct - - pXP496, gaagaaatataagagccaccatggatccgaaggccga + caagccccgcagaaggcagcctacagactgtttacggttggg | This study |
| pXP540 | pVirE – VHH_mCherry__Aat_β_ Spec^R^ | VHH anti mCherry display (cysteine free) (C24A, C98V) on Aat_β_ in pPZP200. Spec^R^   - - pPZP200, AACCGCACCAGGACGG + ccagccagccaacagc - - pXP255, AGCTGTTGGCTGGCTGGCAACGTATCCGCCGTCT + ctggccgtcctggtgcggttctaccacttgatgttgagaccg | This study |
| pXP541 | pVirE – linker_Aat_β_ Spec^R^ | Empty passenger domain (with proline-rich linker) on Aat_β_ in pPZP200. Spec^R^   - - pXP540, gctgaggctcccgatatc + agctgcgtttgcggg - Use of phosphorylated primers and ligation with T4 ligase. | This study |
| pXP542 | pVirE – Aat_β_ Spec^R^ | Empty passenger domain (without proline-rich linker) on Aat_β_ in pPZP200. Spec^R^   - - pXP540, ctctaccgggtcgaggtt + agctgcgtttgcggg - Use of phosphorylated primers and ligation with T4 ligase. | This study |
| pZA002 | pSynth - mScarlet | pGRG36 j23119_mScarlet  Constitutive synthetic promoter driving the expression of mScarlet in Tn7 vector. | Pierrat et. al, 2021 ^18^ |

**Abbreviations:**

Amp = ampicillin

Chloramph = chloramphenicol

gDNA = genomic DNA

Genta = gentamycin

HA tag = YPYDVPDYA = Human influenza hemagglutinin position 98-106

Kan = kanamycin

Neo = neomycin

pCMV = cytomegalovirus promoter

Puro = puromycin

^R^ = resistant

Rif = rifampicin

Spec = Spectinomycin

*Trp1* = phosphoribosylanthranilate isomerase, an enzyme that catalyzes the third step in tryptophan biosynthesis, for auxotrophic selection

**Supplementary table 3: strains used in this study**

| Internal name | Strain name | Description | Reference |
| --- | --- | --- | --- |
| 2 | S17-1 | *E. coli* donor vector for conjugation  pro, res− hsdR17 (rK− mK+) recA− with an integrated RP4-2-Tc::Mu-Km::Tn7, Tpr | Simon *et al.,* 1983^23^ |
| 380 | GV3101 | *Agrobacterium tumefaciens* C58C1 + pMP90, Rif^R^, Genta^R^, Chloramph^R^  The virulence plasmid pC58 was removed from the C58 background and replaced by a disarmed version, pMP90 (Genta^R^) that does not contain a T-DNA sequence. This strain is the starting point for all derivatives. | Koncz *et al*., 1986 ^24^ |
| 464 | *A. tumefaciens* *ΔVirE2* | Markerless *VirE2* knockout generated with pXP233_pNPTS138 in GV3101 | This study |
| 466 | *A. tumefaciens* *VHH_GFP__Aat_β_::ΔVirE2* | VHH anti-GFP cysteine-free display on Aat_β_ was knocked in instead of *VirE2* using pXP234_pNPTS138 in GV3101 | This study |
| 468 | *A. tumefaciens* mScarlet | Constitutive synthetic promoter driving *mScarlet* expression introduced in the *tetR* locus using pXP250 in GV3101 | This study |
| 492 | *S. cerevisiae* eby100 | *Saccharomyces cerevisiae* for yeast display. It produces Aga1 under control of the gal1 promoter and it is trp auxotroph. Parent strain: BJ5465.  Full genomic characteristics: MATa AGA1::GAL1-AGA1::URA3 ura3-52 trp1 leu2-delta200 his3-delta200 pep4::HIS3 prbd1.6R can1 GAL | ATCC mya-4941, gift from B. Correia. |
| AP196 | *S. cerevisiae* eby100 eGFP display | *S. cerevisiae* eby100 containing pXP267 for Aga2p-eGFP display. Selection with the *trp1* auxotrophic marker. | This study |
| Hela tetON GFP disp v4.2 clone 1 | HeLa inducible GFP display  (monoclonal) | Monoclonal cell line of HeLa transduced with lentivectors packaging pXP340, a mammalian lentivector containing a tetracycline-inducible GFP(N105Y, E125V,Y146F) -display anchored with a truncated CD80 transmembrane domain (C-terminal). | Pierrat et. al, 2021 ^18^ |
| HeLa tetON GFP + lenti pXP499 pool puro selected | HeLa GFP inducible display + LgBit (polyclonal) | Polyclonal cell line of HeLa inducible GFP transduced with lentivectors packaging pXP499 - constitutive hPGK promoter driving LgBit (split NanoLuc) expression | This study |
| HEK293T GFPdisp | HEK GFP constitutive (monoclonal) | Monoclonal cell line of HEK293T cells containing pXP145 - pCMV-driven GFP(N105Y, E125V,Y146F) -display anchored with a truncated CD80 transmembrane | This study |
